# Cell-type-specific efferocytosis determines functional plasticity of alveolar macrophages

**DOI:** 10.1101/2023.03.07.528464

**Authors:** Julian Better, Michael Wetstein, Mohammad Estiri, Christina Malainou, Maximiliano Ruben Ferrero, Martin Langelage, Irina Kuznetsova, Ivonne Vazquez-Armendariz, Lucas Kimmig, Siavash Mansouri, Rajkumar Savai, Jochen Wilhelm, Ioannis Alexopoulos, Natascha Sommer, Susanne Herold, Ulrich Matt

## Abstract

Resolution of lung injuries is vital to maintain gas exchange. Concurrently, there is an increased risk of secondary bacterial infections. Alveolar macrophages (AMs) are crucial to clear bacteria and control initiation and resolution of inflammation, but environmental cues that switch functional phenotypes of AMs remain elusive. Here, we discovered an incapacity of AMs to mount an effective immune response to bacteria during resolution of inflammation. AM efferocytosis of neutrophils (PMNs), a hallmark of resolution of inflammation, switched mitochondrial metabolism to shift AM functions. Mechanistically, PMN-derived myeloperoxidase (MPO) fueled canonical glutaminolysis via uncoupling protein 2 (UCP2) resulting in decreased mtROS-dependent killing of bacteria and secretion of pro-inflammatory cytokines. Instead, MPO-enhanced UCP2 expression inhibited mitochondrial hyperpolarization and boosted efferocytosis irrespective of the presence of bacterial pathogens. In contrast, efferocytosis of epithelial cells resulted in a distinct anti-inflammatory phenotype of AMs maintaining phenotypic plasticity towards bacteria. Overall, uptake of apoptotic PMNs switches AMs to prioritize resolution of inflammation over antibacterial responses and similarly affects murine macrophages at extra-pulmonary sites, and human AMs.

**One sentence summary:** Neutrophil efferocytosis reprograms mitochondrial metabolism to switch alveolar macrophages to a pro-resolution phenotype at the cost of bacterial control.

## Introduction

Tissue-resident macrophages (TRMs) regulate inflammation by releasing cytokines or chemokines, clearing pathogens, and promoting resolution of inflammation (e.g., by removal of apoptotic cells) (*1*). Thus, TRMs must swiftly adapt anti- and pro-inflammatory properties with the ultimate goal of restoring homeostasis. In the alveoli and larger airways, AMs are the sentinel cells essential for homeostasis, induction and resolution of inflammation, and clearance of smaller amounts of bacteria to prevent invasive infection (*2, 3*). Importantly, diminished antibacterial properties of AMs facilitate bacterial pneumonia (*4–6*). After lung injuries, restoration of proper gas exchange is vital and requires rapid clearance of apoptotic PMNs by AMs (*7–9*). In macrophages in general, efferocytosis (i.e., the uptake of apoptotic cells by macrophages) has been shown to induce an anti-inflammatory profile characterized by secretion of IL-10 and TGF-β (*10, 11*), enhance clearance of dead cells (*12*), promote organ repair (*13–15*), and decrease antibacterial properties (*16*). Thus, the net benefit of improved resolution might coincide with an impaired ability to fend off bacterial pathogens, which might tip the state of colonization/bacterial control to invasive infection. In line, respiratory viruses, which are leading causes of CAP (*17*), predispose to bacterial superinfections (*18, 19*), and pulmonary aspiration of gastric content was shown to be the strongest independent risk factor for bacterial pneumonia in intubated patients (*20, 21*). Thus, exploring mechanisms that govern antibacterial and pro-resolving functions of AMs is of clinical importance. However, whether these opposing macrophage effector functions are under control of a common upstream signaling hub that regulates the switch of effector functions and the involved cellular compartments and molecular events remained incompletely understood.

We observed earlier that acid aspiration leads to uncontrolled bacterial outgrowth in secondary bacterial pneumonia (*4*). In this model, administration of an anti-inflammatory compound during acid aspiration restored bacterial control by AMs. Thus, inflammation or the resolution thereof affects antibacterial properties of AMs, but the mechanism and meaning of this functional alteration remained unresolved.

Here, we set out to explore functional properties of AMs during inflammation and untangle mechanisms that regulate functional fate decisions of AMs *in vivo*. By combining *in vivo* with functional *ex vivo* as well as *in vitro* analyses, we ultimately revealed that during resolution of inflammation, AMs have impaired antibacterial but improved pro-resolving properties. Mechanistically we found that efferocytosis of neutrophils and not epithelial cells rewires mitochondrial metabolism via MPO in an UCP2-dependent manner, which impaired mtROS production in response to bacteria. Uptake of both apoptotic epithelial cells and neutrophils enhanced efferocytosis; the presence of bacteria favored antibacterial responses after efferocytosis of epithelial cells, while functional plasticity was lost after uptake of apoptotic neutrophils in an MPO-dependent manner. MPO-induced reprogramming of macrophages similarly occurred in viral pneumonia, across bacterial pathogens, and different macrophage subsets, including human AMs (hAMs). Thus, increased UCP2 expression consecutive to PMN efferocytosis represents a conserved immunometabolic decision point that restricts the functional plasticity of TRMs to prioritize resolution of inflammation over host defense against bacteria.

## Results

### AMs exhibit a transient impairment of antibacterial responses along with increased oxidative phosphorylation during resolution of sterile pneumonitis

To explore the functional polarization of AMs during the course of pulmonary inflammation, we used a mouse model of acid aspiration (*4*). This self-resolving model robustly recapitulates key features of lung injury like neutrophil influx, edema formation, and tissue damage (*22*). The major macrophage subsets are tissue-resident AMs, as contribution of monocyte-derived macrophages (MDMs) is marginal and peaks 96 hours after aspiration at low levels (Fig. 1A; gating strategy, fig. S1A). PMN influx is highest around 12 hours after instillation of acid and resolves subsequently (Fig. 1B). To study functional polarization over time, we instilled hydrochloric acid (HCl) intratracheally at different time points, ranging from 12 hours to 8 days. Subsequently, mice were either infected with *Pseudomonas aeruginosa* (*P. aeruginosa*) *in vivo*, or AMs were harvested for further *ex vivo* analysis (Fig. 1C). Bacterial burden was quantified at an early time point (=3 hours) after bacterial infection when *de novo* PMN recruitment is about to begin in sham-treated animals, while a more rapid influx of PMNs was detected in animals 24 hours after acid aspiration (fig. S1B). Despite higher numbers of neutrophils, bacterial outgrowth in bronchoalveolar lavage fluid (BALF) was highest in animals that received acid 24 hours before (Fig. 1D). At this time point of infection, acid pneumonitis is resolving as neutrophils are in decline (Fig. 1B). To investigate if the *in vivo* susceptibility to bacterial infection during resolution of inflammation correlates with AM function, we isolated AMs at the same time points as we infected mice *in vivo* and analyzed their bactericidal properties *ex vivo*. As total AM counts drop during the course of inflammation (Fig. 1A), we normalized AM numbers for *ex vivo* experiments. Matching our *in vivo* data, AMs taken from mice 24 hours after acid aspiration were most significantly impaired in bacterial killing *ex vivo* (Fig. 1E). Thus, our results suggest that, despite the rapid influx of PMNs upon secondary bacterial challenge 24 hours after acid aspiration, a lack and functional defect of AMs account for the increased susceptibility to secondary bacterial infections at this time point.

**Fig. 1.**
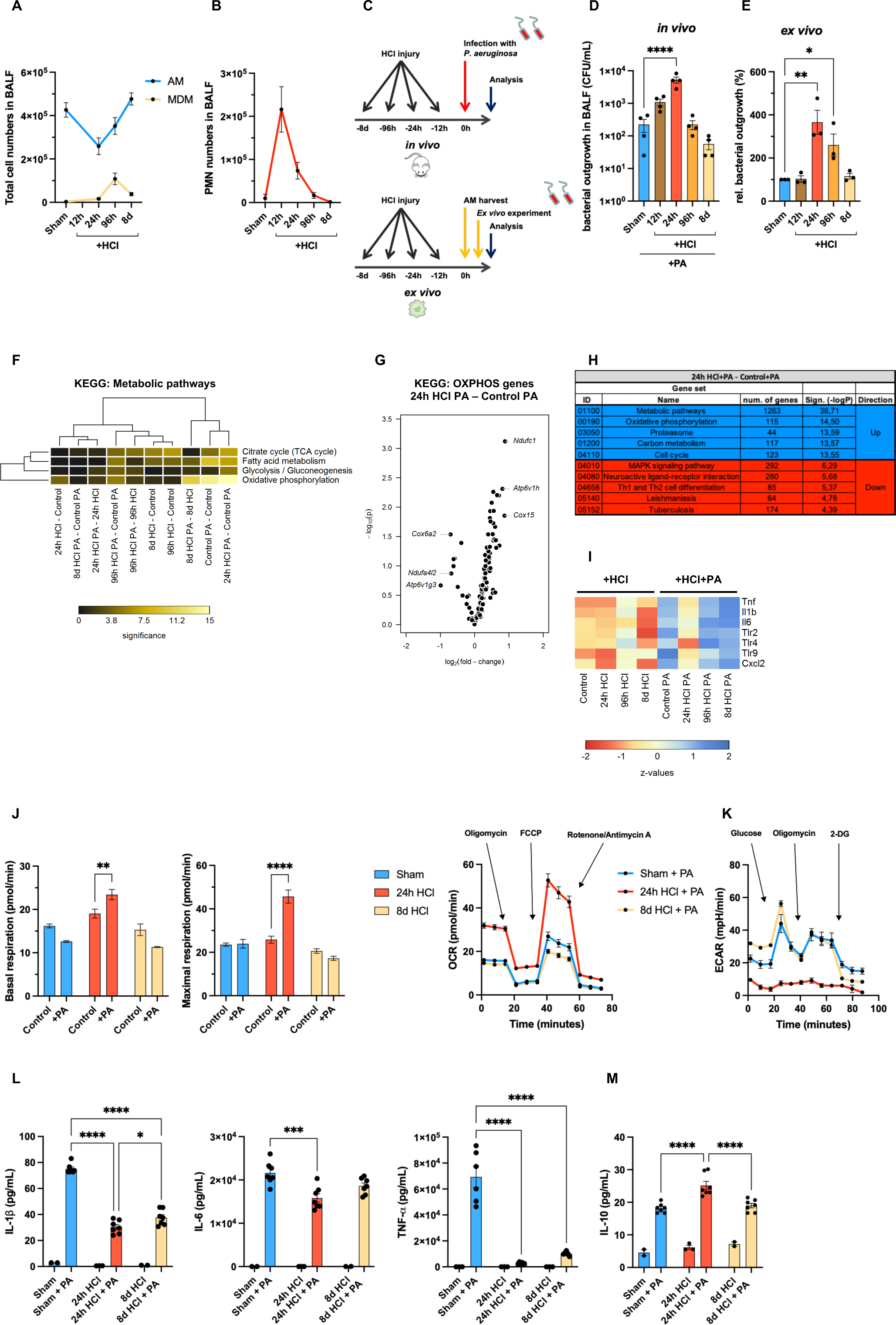
Functional and transcriptional properties of AMs during acid aspiration and secondary bacterial infection. (**A**) AM + MDM numbers in BALF during acid aspiration (without infection) over time (n=7-8 mice/group, pooled analysis from 2-3 independent experiments). (**B**) PMN numbers in BALF during acid aspiration (without infection) over time (n=5-8 mice/group, pooled from 2-3 independent experiments). (**C**) Experimental scheme for *in vivo* and *ex vivo* experiments. (**D**) Bacterial burden in BALF after different time points of acid aspiration and consecutive infection with *P. aeruginosa* (PA) *in vivo* for 3 hours (n=4 mice/group, pooled from two independent experiments). (**E**) Bacterial killing capacity (*P. aeruginosa*) of AMs after different time points of acid aspiration *ex vivo* (n=3 replicates/group, pooled from 3-5 mice/group, representative of 2-3 independent experiments). (**F** to **I**) Bulk mRNA sequencing data of sorted AMs during acid aspiration (24h – 8d HCl) +/− infection with PA (3 hours) presented as KEGG metabolic pathway analysis (F), volcano plot of differentially regulated OXPHOS genes between AMs retrieved from mice 24 hours after acid aspiration + infection with PA compared to infection with PA alone (G), most significantly altered KEGG pathways (H), heat-map of selected pro-inflammatory genes (I) (Bulk mRNA sequencing: n=4 mice/group). (**J**) Oxygen consumption rate (OCR) during Mito Stress Test (extracellular flux assay) of AMs after acid aspiration *ex vivo* in response to *P. aeruginosa* for 6-8 hours; basal respiration (left), maximal respiration (middle) and full analysis for OCR (right) (n=3-6 replicates/group, pooled from 3-5 mice/group, representative of two independent experiments). (**K**) Extracellular acidification rate (ECAR) during Glyco Stress Test (extracellular flux assay) of AMs after acid aspiration *ex vivo* in response to *P. aeruginosa* for 6-8 hours (n=6-16 replicates/group, pooled from 5-8 mice/group and two independent experiments). (**L**) IL-1β (left), IL-6 (middle), and TNF-α (right) secretion of AMs after acid aspiration (24h – 8d HCl) in response to PA for 6-8 hours *ex vivo* (n=3-7 replicates/group, pooled from 3-6 mice/group and two independent experiments). (**M**) IL-10 secretion of AMs after acid aspiration (24h – 8d HCl) in response to PA for 6-8 hours *ex vivo* (n=3-7 replicates/group, pooled from 3-6 mice/group and two independent experiments). *p < 0.05, **p < 0.01, ***p < 0.001, ****p < 0.0001 by one-way ANOVA with Dunnet’s multiple comparisons test (**D** and **E**) or two-way ANOVA with Sidak’s multiple comparisons test (**J**, **L** and **M**). Statistical significance in (**G** and **H**) is presented as negative log p-values. Data are shown as mean ± s.e.m.

Having established the susceptibility to secondary bacterial pneumonia and bactericidal properties of AMs during the course of inflammation, we next performed a transcriptome analysis of flow-sorted AMs after acid aspiration with and without bacterial infection using the same *in vivo* experimental setup (Fig. 1C). Notably, the two most significantly altered pathways in the Kyoto Encyclopedia of Genes and Genomes (KEGG) analysis in all group comparisons performed were “metabolic pathways” (-logP=38,71), followed by “oxidative phosphorylation” (OXPHOS) (-logP=14.5) both between AMs of mice that underwent acid aspiration 24 hours before infection and AMs from animals that were solely infected (Fig. 1F for metabolic pathways and Fig. 1G for OXPHOS genes). Top affected pathways at the vulnerable time point (i.e., 24 hours after acid aspiration) are depicted in Fig. 1H and point towards significant metabolic rewiring, an increase in cell cycle and proteasome transcripts, while immune responses are downregulated. Accordingly, analysis of selected pro-inflammatory marker genes such as *Tnf-α*, *Il-6*, *Il-1β*, *Cxcl2*, and Toll-like receptors (*tlr*s) illustrates an impaired response to bacteria 24 hours after acid aspiration in AMs (Fig. 1I), which was similarly found in KEGG-pathway analysis of NFκB-, TNF-, and mitogen-activated protein (Map)-kinase signaling (Fig. S1, C to E). At later time points after instillation of acid, a pro-inflammatory response recurred (Fig. 1I). Thus, 24 hours after acid aspiration AMs are less responsive to a bacterial challenge. To test if transcriptional metabolic changes mirror functional properties, we performed *ex vivo* extracellular flux analyses. Basal and maximal respiration were increased in response to bacteria in AMs of animals that underwent acid aspiration 24 hours before compared to sham-treated animals and those that received acid 8 days before (Fig. 1J), linking transcriptional regulation to mitochondrial function. At the same time, AMs retrieved 24 hours after acid aspiration showed a clearly diminished extracellular acidification rate (ECAR) when challenged with bacteria compared to sham treatment (Fig. 1K). Concurrently, secretion of pro-inflammatory cytokines (IL-1β, IL-6, and TNF-α) was significantly decreased in AMs of mice during resolution of inflammation compared to sham-treated animals *ex vivo* (Fig. 1L). Vice versa, the anti-inflammatory cytokine IL-10 was highest upon stimulation with *P. aeruginosa* when acid had been applied 24 hours before (Fig. 1M).

Thus, AMs exhibit a transiently impaired response to bacteria, paralleled by an increased oxygen consumption rate (OCR).

### Production of mtROS is impaired during resolution of inflammation in AMs upon bacterial encounter

During OXPHOS, mitochondria produce mtROS (*23*), which is a crucial process in bacterial killing (*24, 25*). Therefore, we tested the capacity of acid-experienced versus sham-treated AMs to produce mtROS in response to *P. aeruginosa ex vivo*. AMs harvested 24 hours after acid aspiration failed to mount mtROS production upon bacterial stimulation, whereas, after 8 days, mtROS release recurred (Fig. 2A and fig. S2A). In contrast, cytosolic ROS production (cROS) remained intact in AMs of mice that underwent acid aspiration (Fig. 2B). Flux through the electron transport chain (ETC) gives rise to a basal amount of mtROS release. An increased mitochondrial membrane potential (MMP) results in enhanced mtROS production and may limit electron flux, oxygen consumption, and ATP production under certain conditions (*26*). As such, the MMP was shown to be a driving force for the generation of mtROS (*27*), and to be required for mtROS release in response to LPS (*28*). Accordingly, AMs from mice that underwent acid aspiration 24 hours before failed to increase the MMP upon bacterial stimulation *ex vivo*, whereas AMs from sham-treated animals or from mice that underwent aspiration 8 days before responded with an increase of the MMP (Fig. 2C). Confirming the importance of mtROS for bacterial control, the mtROS scavenger MitoTEMPO reduced intracellular bacterial killing in AMs from sham-treated animals and from mice that had received acid 8 days or 12 hours before but did not alter bacterial clearing capacity of AMs 24 hours after acid aspiration *ex vivo* (Fig. 2D). Apart from affecting bactericidal properties, mtROS were shown to impact the secretion of cytokines(*29*). Scavenging of mtROS in AMs *in vitro* markedly diminished secretion of IL-1β and IL-6 in response to *P. aeruginosa* (Fig. 2E). However, release of TNF-α and the anti-inflammatory cytokine IL-10 remained unaffected in this setting (Fig. 2, E and F).

**Fig. 2.**
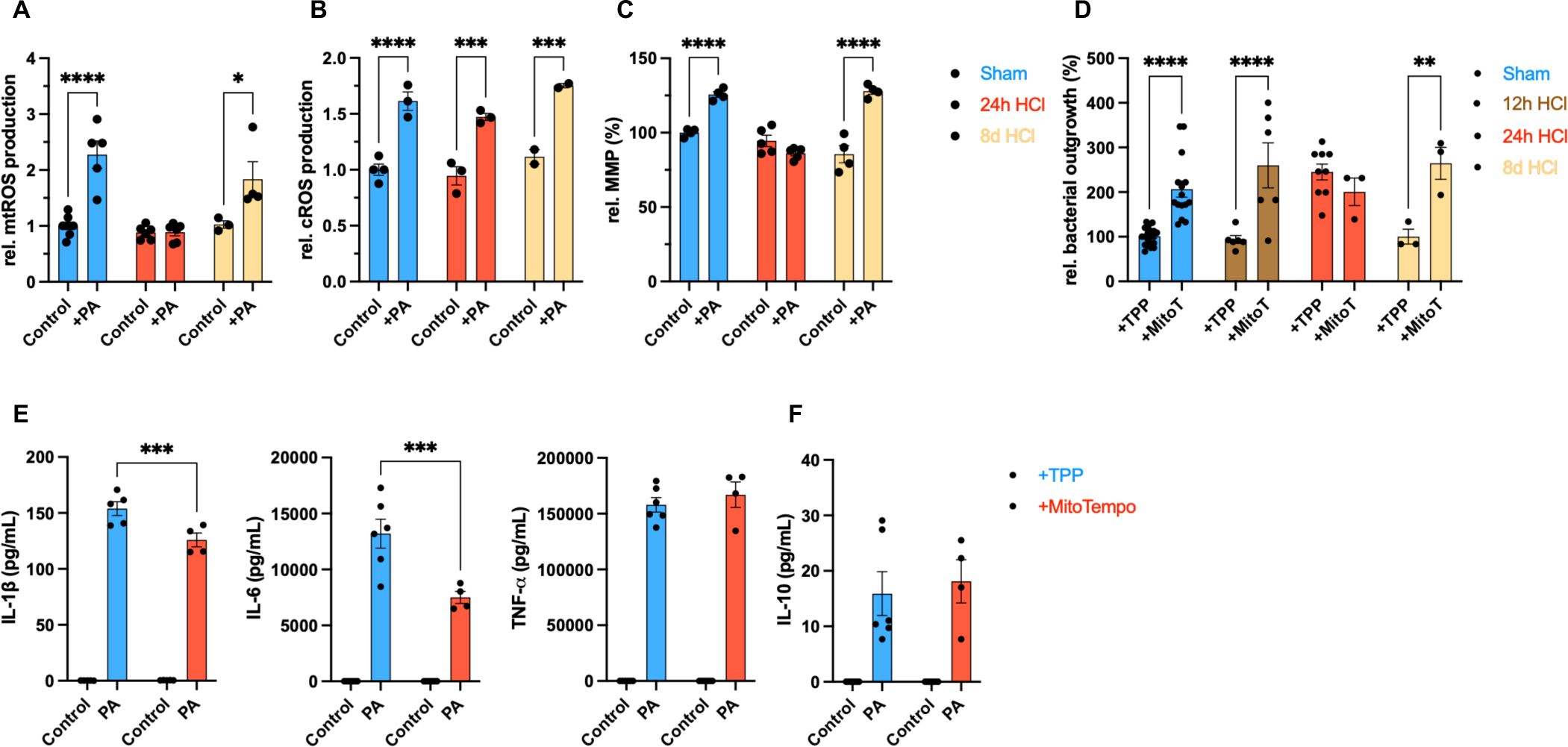
AMs fail to mount mtROS in response to bacteria after acid aspiration. (**A** to **D**) *Ex vivo* analyses of AMs after acid aspiration at indicated time-points for mtROS (A), cROS (B), and MMP (C) in response to PA for 6-8 hours (n=3-6 replicates/group, pooled from 5-8 mice/group, representative of 2-3 independent experiments); (D) Bacterial killing of *P. aeruginosa* +/− MitoTEMPO (MitoT) 100μm for 1 hour after acid aspiration *ex vivo* (Control/Carrier: TPP) (n=3-21 replicates/group, pooled from 3-12 mice/group and 2-3 independent experiments. (**E**) IL-1β (left), IL-6 (middle), and TNF-α (right) secretion of AMs +/− MitoTEMPO 100μm for 1 hour after acid aspiration *ex vivo* (control/carrier: TPP) in response to PA for 6-8 hours (n>=3-6 replicates/group, pooled from 3-5 mice/group, representative two independent experiments). (**F**) IL-10 secretion of AMs +/− MitoTEMPO 100μm for 1 hour after acid aspiration *ex vivo* (control/carrier: TPP) in response to PA for 6-8 hours (n>=3-6 replicates/group, pooled from 3-5 mice/group, representative two independent experiments). *p < 0.05, **p < 0.01, ***p < 0.001, ****p < 0.0001 by two-way ANOVA with Sidak’s multiple comparisons test (**A** to **F**). Data are shown as mean ± s.e.m.

Thus, AMs have a profound defect in generating mtROS in response to bacteria during resolution of inflammation affecting mtROS-dependent bacterial killing and pro-inflammatory cytokines.

### Efferocytosis of PMNs leads to increased oxygen consumption and precludes mtROS release in response to bacteria in AMs

Next, we aimed to delineate which alveolar cues switch AMs towards an impaired bacterial response during resolution of acid-induced pneumonitis. Incubation of BALF from mice after acid aspiration did not alter bactericidal properties of AMs (fig. S3A), rendering soluble mediators less likely to account for impaired bacterial defense. However, cellular composition of BALF after acid aspiration revealed increased levels of apoptotic epithelial cells (main constituent of CD45neg cells) and apoptotic PMNs peaking at around 12 to 24 hours (Fig. 3A). Since AMs are important for clearance of apoptotic cells during resolution of inflammation, we speculated that efferocytosis might affect innate immune effector functions in this context. To assess their potential impact on mtROS production, apoptosis in alveolar epithelial cells (AECs) and PMNs was induced with staurosporine to similar levels (fig. S3B). Whereas uptake of apoptotic AECs by AMs did not affect the release of mtROS, ingestion of apoptotic PMNs abolished mtROS production upon bacterial stimulation with *P. aeruginosa* and *S. pneumoniae in vitro* (Fig. 3B and fig. S3C, respectively). Accordingly, AM efferocytosis of PMNs, but not of AECs, prevented an increase of the MMP (Fig. 3C) in response to and diminished bacterial killing of *P. aeruginosa* (Fig. 3D). Matching the *ex vivo* data (Fig. 2D), scavenging of mtROS did not further reduce the bacterial killing capacity of AMs after ingestion of apoptotic PMNs *in vitro* (Fig. 3E). Of note, ingestion of apoptotic PMNs had no impact on cROS release upon bacterial encounter (fig. S3D). In line with reduced mtROS generation, cytokine release by AMs was completely (IL-1β) or partially (IL-6) diminished after PMN efferocytosis and consecutive bacterial challenge (Fig. 3F). Uptake of apoptotic AECs slightly reduced IL-6 levels, whereas uptake of either apoptotic AECs or PMNs blunted TNF-α release in response to *P. aeruginosa* (Fig. 3F). Of note, significant baseline IL-10 secretion was detected after efferocytosis of AECs and PMNs by AMs without prior bacterial stimulation (fig. S3E). In contrast, *P. aeruginosa*-treated AMs released IL-10 only consecutive to PMN efferocytosis (Fig. 3G).

**Fig. 3.**
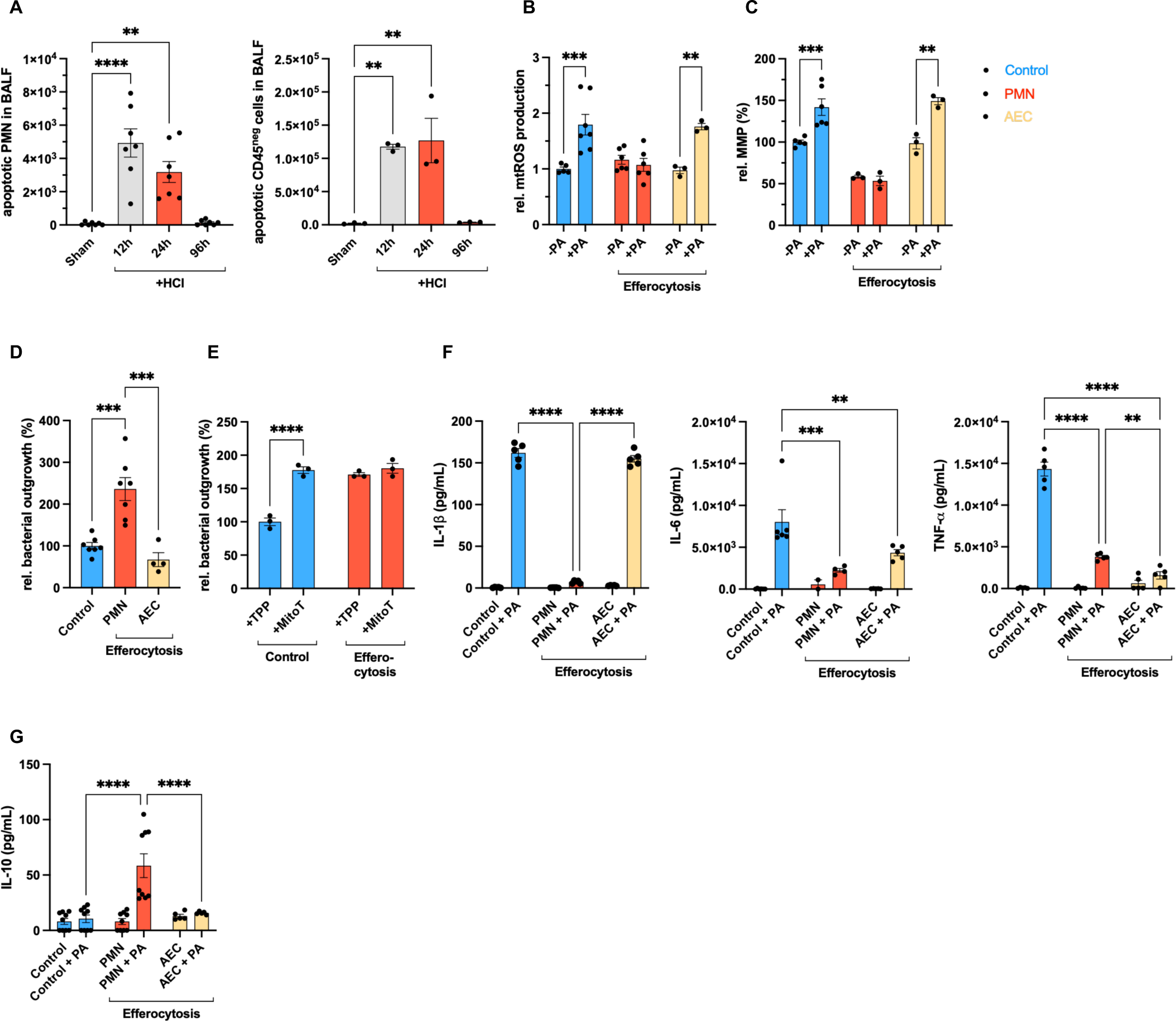
Cell-type specific efferocytosis of PMNs precludes mtROS generation in response to bacteria in AMs. (**A**) Total number of apoptotic PMNs (left) and CD45^neg^ cells (right) after different time points of acid aspiration (12-96h HCl) in the BALF (n=3-7 mice/group, pooled from two independent experiments). (**B**) mtROS release of AMs after efferocytosis of apoptotic PMNs or alveolar epithelial cells (AECs) for 4 hours in response to PA (n=3-7 replicates/group, pooled from 3-5 mice/group and two independent experiments). (**C**) MMP measurement in AMs after efferocytosis of apoptotic PMNs or AECs for 4 hours in response to PA (n=3-6 replicates/group, pooled from 3-5 mice/group and two independent experiments). (**D**) Bacterial killing capacity (*P. aeruginosa*) of AMs after efferocytosis of apoptotic PMNs or AECs for 4 hours (n=3-7 replicates/group, pooled analysis from 3-5 mice/group and two independent experiments). (**E**) Bacterial killing capacity (*P. aeruginosa*) of AMs after efferocytosis of apoptotic PMNs compared to control (no efferocytosis) +/− MitoT 100μm for 1 hour (control/carrier: TPP) (n=3 replicates/group, pooled from 4 mice/group and two independent experiments). (**F**) IL-1β (left), IL-6 (middle), and TNF-α (right) secretion of AMs after efferocytosis of apoptotic PMNs or AECs for 4 hours in response to PA for 6-8 hours (n=3-6 replicates/group, pooled from 3-5 mice/group and two independent experiments). (**G**) IL-10 secretion of AMs after efferocytosis of apoptotic PMNs or AECs for 4 hours in response to PA for 6-8 hours (n=5-9 replicates/group, pooled from 4-7 mice/group and two independent experiments). *p < 0.05, **p < 0.01, ***p < 0.001, ****p < 0.0001 by one-way ANOVA with Dunnet’s (**A**) or Tukey’s multi comparisons (**D**) test or two-way ANOVA with Sidak’s multiple comparisons test (**B** and **C**, **E** to **G**). Data are shown as mean ± s.e.m.

Taken together, our data demonstrate that the effects of efferocytosis on the functional phenotype of AMs are cell-type specific. PMN but not AEC ingestion blunted mtROS release in AMs during bacterial challenge, which ultimately impaired bacterial killing and diminished secretion of pro-inflammatory cytokines, while IL-10 secretion was enhanced.

### PMN-derived MPO mediates immunometabolic alterations upon efferocytosis in AMs in response to bacteria

We next sought to identify the PMN component accounting for efferocytosis-driven immunometabolic alterations in AMs. As enzymes are major constituents of PMN granules mediating antimicrobial functions, we tested lipocalin-2 (NGAL), elastase, and MPO for their ability to inhibit mtROS production upon bacterial stimulation in AMs. Whereas elastase and NGAL-treated AMs maintained their capacity to mount mtROS, pre-incubation with MPO abrogated mtROS production in response to *P. aeruginosa* (Fig. 4A). Of note, the concentration of MPO (0.5µM) used did not affect the viability of AMs (fig. S4A). Mitochondrial content was neither altered by MPO treatment *in vitro* (Fig. 4B) nor 24 hours after acid aspiration *ex vivo* (fig. S4B). To elucidate MPO᾽s impact in the context of efferocytosis, we incubated AMs with apoptotic PMNs of either wildtype (WT) or *Mpo*^−/−^ animals. MPO deficiency clearly restored the capacity of AMs to build mtROS (Fig. 4C) and led to an increase of the MMP after efferocytosis in response to bacteria (Fig. 4D). Accordingly, the killing capacity of AMs improved significantly after incubation with PMNs lacking MPO compared to WT PMNs (Fig. 4E). Extracellular flux analysis of AMs that had ingested *Mpo*^−/−^ PMNs showed lower basal and maximal respiration compared to WT PMNs when challenged with bacteria, further confirming that heightened OXPHOS correlates with a dampened antibacterial response (Fig. 4F). To address the relevance of our findings *in vivo*, we instilled apoptotic PMNs intratracheally and subsequently infected mice with *P. aeruginosa* (Fig. 4G). Consistent with our *in vitro* results, a lack of MPO in PMNs improved bacterial clearance compared to WT PMNs *in vivo* (Fig. 4H). Of note, PMN efferocytosis partially diminished bacterial killing *in vitro* and clearance *in vivo* independent of MPO (Fig. 4E and Fig. 4H, respectively). Thus, as previously reported, additional mechanisms such as prostaglandin E-2 mediated inhibition of bacterial killing most likely play a role (*16*). To examine if MPO attenuation of bacterial control is a pathogen-specific effect, we tested *Klebsiella pneumoniae* (*K. pneumoniae*) and *S. pneumoniae* in this experimental setting. Elimination of both pathogens was diminished by pre-incubation with MPO *in vitro* (Fig. 4I). Taken together, PMN-derived MPO, likely unleashed during the efferocytosis process, abrogates mtROS production and increases oxygen consumption in response to bacteria in AMs.

**Fig. 4.**
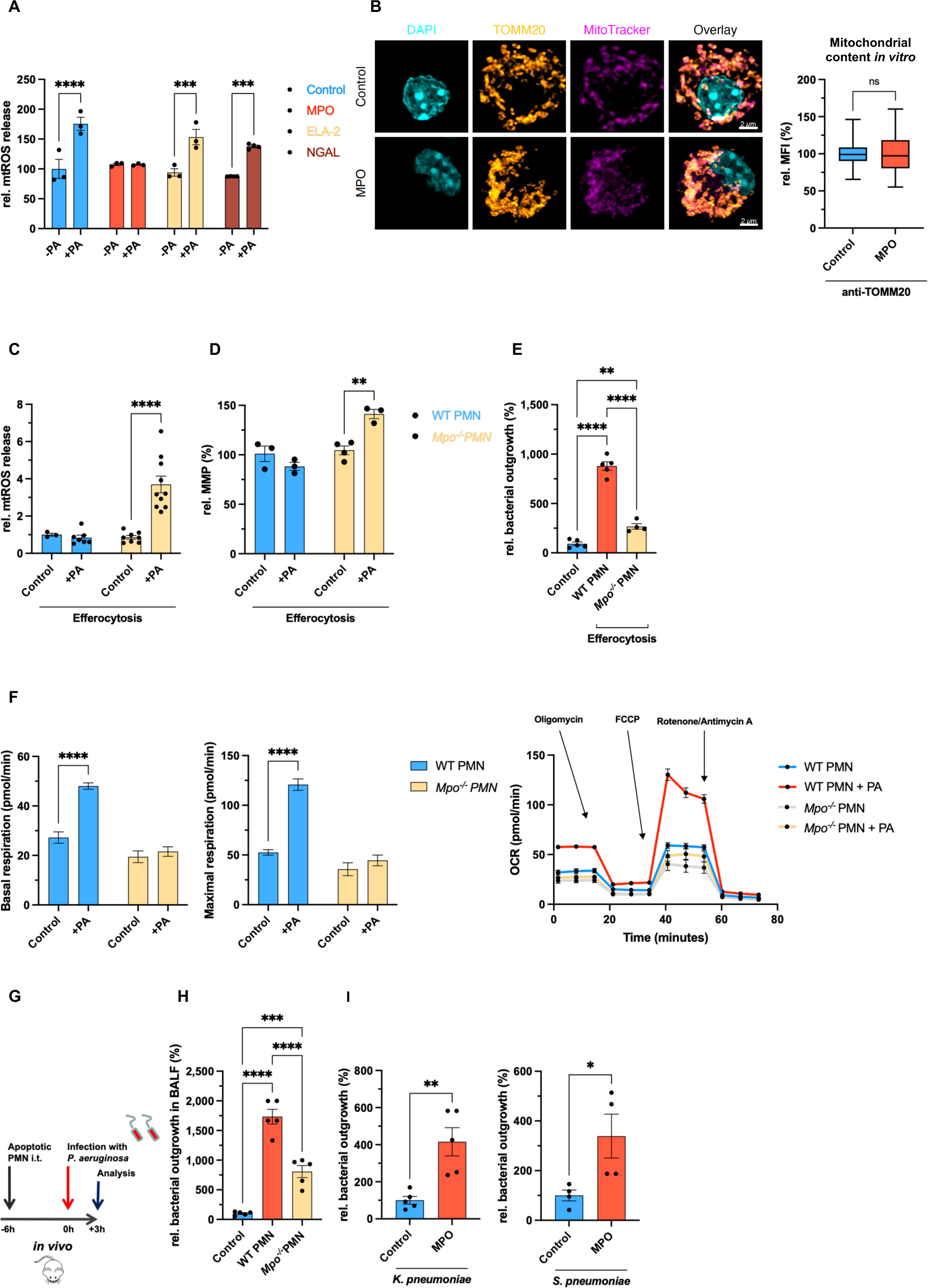
PMN-derived MPO mediates alterations of AM functions in response to bacteria. (**A**) mtROS measurement in AMs after incubation with MPO, ELA-2, or NGAL each 0.5µM for 3 hours in response to PA (n=3-4 replicates/group, pooled from three mice/group and two independent experiments). (**B**) Confocal microscopy of AMs after treatment with MPO; nuclei were stained with DAPI, mitochondria with anti-translocase of outer mitochondrial membrane 20 (TOMM20) and MitoTracker deep red (left); quantification of the mean fluorescence intensity of anti-TOMM20 normalized to control reflecting mitochondrial content *in vitro* (right) (n=4 mice/group, pooled from two independent experiments). (**C**) mtROS measurement in AMs after efferocytosis of apoptotic PMNs of wild-type (WT) or Myeloperoxidase-deficient (*Mpo*^−/−^) mice for 4 hours in response to PA for 6-8 hours (n=3-10 replicates/group, pooled from 3-8 mice/group and two independent experiments). (**D**) MMP measurement in AMs after efferocytosis of apoptotic PMNs of WT or *Mpo*^−/−^ mice for 4 hours in response to PA for 6-8 hours (n=3-4 replicates/group, pooled from 3-4 mice/group and two independent experiments). **(E)** Bacterial killing capacity (*P. aeruginosa*) of AMs after efferocytosis of PMNs of WT or *Mpo*^−/−^ mice (n=4-5 replicates/group, pooled from four mice/group and two independent experiments). (**F**) OCR during Mito Stress Test (extracellular flux assay) of AMs after efferocytosis of apoptotic PMNs of WT or *Mpo*^−/−^ mice for 4 hours in response to PA for 6-8 hours; basal respiration (left), maximal respiration (middle) and full analysis for OCR (right) (n=3-6 replicates/group, pooled from 3-5 mice/group and two independent experiments). (**G**) Experimental scheme for (H). (**H**) Bacterial outgrowth in BALF after instillation of apoptotic PMNs of WT or *Mpo*^−/−^ mice and subsequent infection with *P. aeruginosa* for 3 hours (n=5 mice/group, pooled from two independent experiments). (**I**) Bacterial killing capacity (left: *K.* pneumoniae; right: *S. pneumoniae*) of AMs +/− pre-treatment with MPO 0.5μM for 3 hours (n=4-5 replicates/group, pooled from four mice/group and two experiments). *p < 0.05, **p < 0.01, ***p < 0.001, ****p < 0.0001 by Student’s t-test (**B**, **I**) or one-way ANOVA with Tukey’s multi comparisons test (**E**, **H**) or two-way ANOVA with Sidak’s multiple comparisons test (**A**, **C** and **D**, **F**). Data are shown as mean ± s.e.m..

### MPO increases UCP2 expression, which mediates inhibition of mtROS generation upon bacterial encounter

Clearance of apoptotic PMNs rendered AMs unable to increase the MMP and consecutively the release of mtROS upon bacterial encounter while promoting mitochondrial respiration. Uncoupling proteins (UCPs) were found to dissipate the proton gradient across the inner mitochondrial membrane resulting in decreased MMP and mtROS production (*26, 30*). To test if UCP2, which is expressed in macrophages (*31*) and AMs (*32*), is implicated in MPO-induced mitochondrial reprogramming, we applied the UCP2 inhibitor genipin. After MPO pre-treatment, genipin rescued mtROS production in AMs stimulated with *P. aeruginosa* and *K. pneumoniae in vitro* (Fig. 5A and fig. S5A, respectively). In line, incubation with genipin restored mtROS release of AMs isolated 24 hours after acid aspiration and challenged with bacteria *ex vivo* (Fig. 5B). Next, we used *Ucp2*^−/−^ AMs, exposed them to apoptotic WT PMNs for efferocytosis *in vitro* and found them to still respond with mtROS production when challenged with bacteria (Fig. 5C). Accordingly, in *Ucp2*^−/−^ AMs, MPO pre-treatment did not preclude an increase of the MMP in response to bacteria (Fig. 5D) and did not diminish bacterial killing *in vitro* (Fig. 5E). Furthermore, the OCR did not rise in response to bacteria after MPO pre-treatment in UCP2-deficient AMs (Fig. 5F). Regarding cytokine secretion, inhibition of UCP2 using genipin restored the release of IL-1β and IL-6, but not TNFα in AMs of WT mice pretreated with MPO *in vitro* (fig. S5B). In contrast, secretion of IL-10 was not affected (fig. S5C). Thus, MPO-mediated impaired mtROS production in AMs in response to bacteria critically depends on UCP2 function. Therefore, we speculated that MPO might affect UCP2 levels in AMs. While we did not detect transcriptional upregulation of UCP2 during acid-induced inflammation *in vivo* or efferocytosis *in vitro* (fig. S5D), protein levels of UCP2 were increased in AMs harvested 24 hours after acid aspiration compared to sham-treated animals (Fig. 5G). In line, *in vitro* stimulation of AMs with MPO increased the expression of UCP2 (Fig. 5H). To demonstrate the relevance of our findings *in vivo*, we treated mice 24 hours after acid aspiration with genipin or carrier control intratracheally and subsequently infected them with *P. aeruginosa* (Fig. 5I). Confirming our *ex vivo* and *in vitro* data, mice that received genipin exhibited a significantly lower bacterial burden in BALF compared to control animals (Fig. 5J). Of note, genipin did neither impair bacterial growth itself *in vitro* nor in mice infected with *P. aeruginosa* without prior acid aspiration *in vivo* (fig. S5E).

**Fig. 5.**
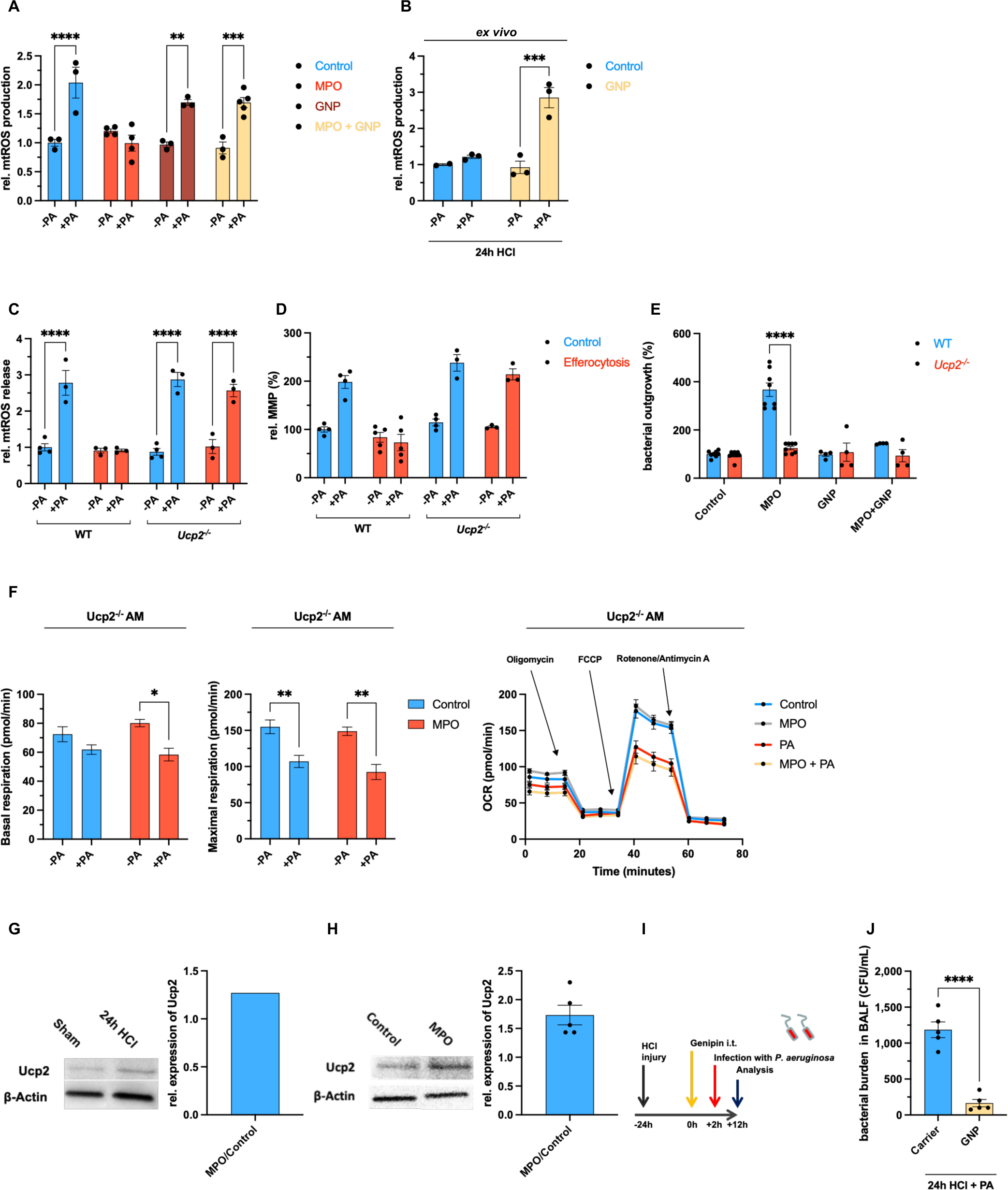
MPO-mediated increase of UCP2 precludes mtROS release in response to bacteria. (**A**) mtROS measurement in AMs pre-treated with MPO 0.5μM for 3 hours +/− genipin (GNP) 100µM for 1 hour in response to PA for 6-8 hours (n=3-5 replicates/group, pooled from 3-4 mice/group and two independent experiments). (**B**) *Ex vivo* mtROS measurement in AMs 24 hours after acid aspiration pre-treated with GNP 100µM for 1 hour compared to control (DMSO) in response to PA for 6-8 hours (n=3 replicates/group, pooled from five mice/group and two independent experiments). (**C**) mtROS measurement in AMs of WT or *Ucp2*^−/−^ AMs after efferocytosis of apoptotic PMNs for 4 hours compared to control (no efferocytosis) in response to PA for 6-8 hours (n=3-4 replicates/group, pooled from three mice/group and two independent experiments). (**D**) MMP measurement AMs of WT or *Ucp2*^−/−^ AMs after efferocytosis of apoptotic PMNs for 4 hours compared to control (no efferocytosis) in response to PA for 6-8 hours (n=3-5 replicates/group, pooled from 3-4 mice/group and two independent experiments). (**E**) Bacterial killing capacity (*P. aeruginosa*) of WT or *Ucp2*^−/−^ AMs pre-treated with MPO 0.5μM for 3 hours +/− GNP 100µM for 1 hour (n=4-8 replicates/group, pooled from 3-6 mice/group and two independent experiments). (**F**) OCR during Mito Stress Test (extracellular flux assay) of *Ucp2*^−/−^ AMs pre-treated with MPO 0.5μM for 3 hours in response to PA for 6-8 hours; basal respiration (left), maximal respiration (middle) and full analysis for OCR (right) (n=4-9 replicates/group, pooled from 4-7 mice/group and two independent experiments). (**G**) Western Blot of Ucp2 and β-Actin (loading control) in AMs 24 hours after acid aspiration compared to sham (whole cell lysate) (pooled from 8-12 mice/group). (**H**) Western Blot of Ucp2 and β-Actin (loading control) in AMs treated with MPO 0.5μM for 3 hours compared to control (n=5 replicates/group, pooled from twenty mice/group and two independent experiments). (**I**) Experimental scheme for (J). (**J**) Bacterial burden in BALF 24 hours after acid aspiration and successive intratracheal instillation of GNP 300µM for 2 hours compared to control (DMSO) followed by *in vivo* infection with PA for 10 hours (n=5 mice/group, pooled from two independent experiments). *p < 0.05, **p < 0.01, ***p < 0.001, ****p < 0.0001 by Student’s t-test (**J**) or two-way ANOVA with Sidak’s multiple comparisons test (**A** and **B**, **E** and **F**) or three-way ANOVA with Sidak’s multiple comparisons test (**C** and **D**). Data are shown as mean ± s.e.m.

Together, these data reveal that PMN-derived MPO increases UCP2 expression in AMs, augmenting oxygen consumption during bacterial stimulation while impairing infection-induced mtROS production and consecutively bacterial killing and cytokine secretion.

### MPO-induced UCP2 expression drives canonical glutaminolysis in response to bacteria

Emerging evidence highlights the role of UCP2 as a metabolic hub regulating the relative contribution of substrates oxidized by mitochondria (*33–36*). More precisely, UCP2 favors the oxidation of fatty acids and glutamine while limiting the oxidation of glucose-derived pyruvate (Fig. 6A). Since MPO pre-treatment led to an increase of the OCR via UCP2 in response to bacteria, we wondered whether specific fueling of the tricarboxylic acid (TCA) cycle is implicated herein. Investigating the maximum mitochondrial respiration in presence of pathway-specific inhibitors (substrate oxidation stress test) revealed a relative increase in glutaminolysis and conversely, a relative decrease in glycolysis in AMs when treated with *P. aeruginosa* plus MPO compared to *P. aeruginosa* alone, whereas fatty acid oxidation remained unchanged (Fig. 6B). In fact, inhibition of glutaminolysis by BPTES, an inhibitor of glutaminase-1, completely blunted an increase in maximal respiration in response to MPO and bacteria *in vitro* (Fig. 6C). Furthermore, wildtype AMs that were pre-treated with MPO and stimulated with bacteria showed increased glutamine consumption and intracellular levels of glutamate while intracellular levels of glutamine remained unaffected compared to control (Fig. 6D). In addition, MPO pre-treated AMs showed diminished lactate release and glucose consumption in presence of bacteria (Fig. S6, A and B). These MPO-mediated metabolic alterations were absent in UCP2-deficient AMs (Fig. 6D and Fig. S6, A and B). Accordingly, inhibition of glutaminolysis reversed the MPO-mediated impairment in mtROS production during bacterial encounter (Fig. 6E). In line, the inhibitory effect of PMN efferocytosis on the release of the mtROS-dependent cytokine IL-1β in response to bacteria was partially reversed after inhibition of glutaminolysis (Fig. 6F), while, conversely, IL-10 secretion was reduced (Fig. 6G). Canonical glutaminolysis relies on the enzymatic activity of glutamate dehydrogenase (GLUD1), which converts glutamate into α-ketoglutarate (α-KG) to fuel the TCA cycle. Treatment of AMs with the GLUD1-inhibitor epigallocatechin gallate (EGCG) after pre-treatment with MPO restored mtROS production (fig. S6C), and the increase of the MMP (fig. S6D) in response to *P. aeruginosa*. Dimethyl-α-KG (DM-α-KG) is a cell-permeable analog of α-KG. Treatment of AMs with genipin plus DM-α-KG restored the MPO-mediated inhibition of mtROS secretion (Fig. 6H), enabled an increase of the MMP in response to bacteria (Fig. 6I), and normalized the capacity to kill bacteria (Fig. 6J).

**Fig. 6.**
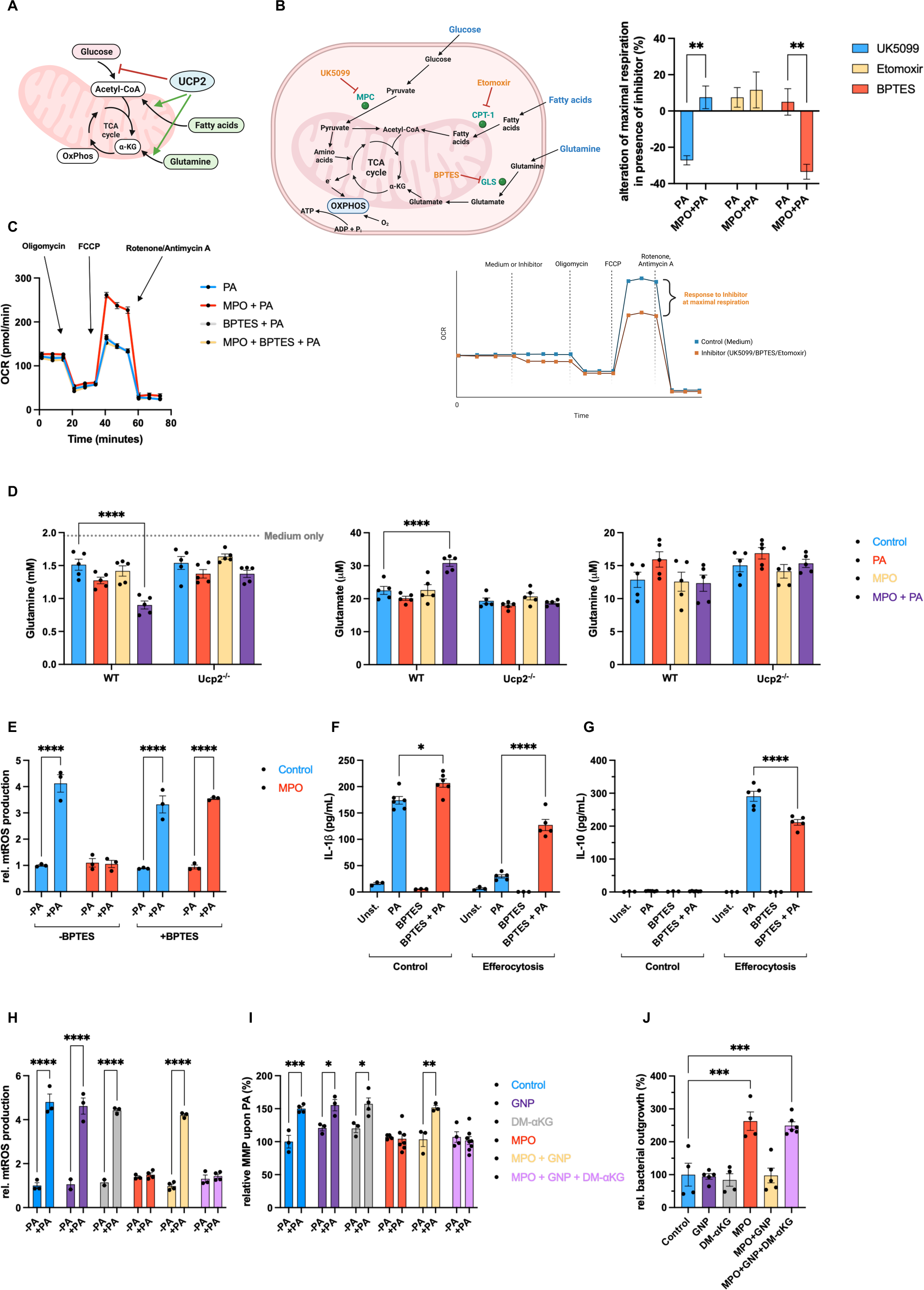
UCP2-mediated alterations of AMs in response to bacteria depend on glutaminolysis. (**A**) Diagram of UCP2 mediated rewiring of mitochondrial metabolism. (**B**) Substrate Oxidation Stress test (extracellular flux assay): Primary metabolic pathways, including glycolysis, fatty acid oxidation, glutaminolysis, TCA cycle, and OXPHOS; pathway-specific inhibitors are highlighted in orange, corresponding enzymes in green (MPC = Mitochondrial Pyruvate Carrier; CPT-1 = Carnitine Palmitoyltransferase 1; GLS = Glutaminase), pathway substrates in blue (top left); exemplary Substrate Oxidation Stress Test (bottom left); data displayed as alteration of maximal respiration in presence of pathway-specific inhibitors (UK5099 for glycolysis, etomoxir for fatty acid oxidation, BPTES for glutaminolysis) compared to respective uninhibited control in AMs pre-treated with MPO 0.5μM for 3 hours compared to control in presence of PA for 6-8 hours (right) (n=7-9 replicates/group, pooled from 6-8 mice/group and two independent experiments). (**C**) OCR during Mito Stress Test (extracellular flux assay) of AMs after pre-treatment with MPO 0.5μM for 3 hours followed by BPTES 3 µM for 1 hour compared to control (DMSO) in response to PA for 6-8 hours (n=5 replicates/group, pooled from four mice/group and two independent experiments). (**D**) Luminometric assessment of glutamine consumption by and intracellular concentration of glutamate and glutamine in AMs from WT or Ucp2^−/−^ mice after pre-treatment with MPO 0.5μM for 3 hours in response to PA for 6-8 hours (n=5 replicates/group, pooled from 4 mice/group and two independent experiments). (**E**) mtROS measurement in AMs after pre-treatment with MPO 0.5μM for 3 hours followed by BPTES 3 µM for 1 hour compared to control (DMSO) in response to PA for 6-8 hours (n=3 replicates/group, pooled from three mice/group and two independent experiments). (**F**) IL-1β secretion of AMs after efferocytosis of apoptotic PMNs for 4 hours +/− BPTES in response to PA for 6-8 hours (n=3-6 replicates/group, pooled from 3-5 mice/group and two independent experiments). (**G**) IL-10 secretion of AMs after efferocytosis of apoptotic PMNs for 4 hours +/− BPTES in response to PA for 6-8 hours (n=3-5 replicates/group, pooled from 3-5 mice/group and two independent experiments). (**H**) mtROS measurement in AMs pre-treated with MPO 0.5μM for 3 hours +/− GNP 100μM for 1 hour concomitantly +/− DM-α-KG (αKG) 1mM for 1 hour in response to PA (n=3-4 replicates/group, pooled from four mice/group and two independent experiments). (**I**) MMP measurement in AMs pre-treated with MPO 0.5μM for 3 hours +/− GNP 100μM for 1 hour concomitantly +/− DM-α-KG (αKG) 1mM for 1 hour in response to PA (n=3-7 replicates/group, pooled from 4-7 mice/group and two independent experiments). (**J**) Bacterial killing capacity (*P. aeruginosa*) of AMs pre-treated with MPO 0.5μM for 3 hours +/− GNP 100μM for 1 hour concomitantly +/− DM-α-KG (αKG) 1mM for 1 hour in response to PA for 6-8 hours (n=3-6 replicates/group, pooled from 4-6 mice/group and two independent experiments). *p < 0.05, **p < 0.01, ***p < 0.001, ****p < 0.0001 by one-way ANOVA with Dunnet’s multi comparisons test (**J**) or two-way ANOVA with Sidak’s multiple comparisons test (**B**, **H** and **I**) or three-way ANOVA with Sidak’s multiple comparisons test (**D** to **G**). Data are shown as mean ± s.e.m.

Together, MPO-induced UCP2 expression enhances the efficient oxidation of glutamine via canonical glutaminolysis in the presence of bacteria. Consequently, AMs are unable to mount their mtROS production, which impairs bacterial killing and affects mtROS-dependent cytokine release.

### MPO primes different macrophage subsets to blunt bacterial response

Next, we wanted to investigate if MPO-induced immunometabolic alterations are restricted to AMs or also apply to macrophages of extra-pulmonary sites. As in AMs, efferocytosis of PMNs but not AECs by primary peritoneal macrophages (PMs) and bone marrow-derived macrophages (BMDMs) resulted in decreased mtROS production (Fig. 7A), an unaltered MMP (fig. S7A) in response to, and an impaired killing of *P. aeruginosa* (Fig. 7B), while cROS secretion remained unaffected *in vitro* (fig. S7C). In line, MPO increased UCP2 expression in both macrophage subsets (fig. S7C), and metabolic analysis demonstrated an increased relative contribution of glutaminolysis to mitochondrial oxygen consumption after MPO pre-treatment in response to bacteria in both PMs and BMDMs (fig. S7D). In contrast, the contribution of glycolysis- and fatty acid oxidation to mitochondrial oxygen consumption remained unaltered by MPO treatment (fig. S7D). Furthermore, mtROS production was unaffected after MPO incubation and challenge with *P. aeruginosa* in *Ucp2*^−/−^ BMDMs and PMs (Fig. 7C). Neutrophil efferocytosis also drastically diminished IL-1β secretion in PMs and BMDMs stimulated with bacteria (fig. S7E), suggesting a conserved mechanism among macrophages of different ontogeny and tissues. As in AMs, IL-6 secretion was inhibited after PMN efferocytosis and upon bacterial stimulation in PMs and was unspecifically diminished by efferocytosis in general in BMDMs (fig. S7E). IL-10 secretion was detected already after bacterial stimulation alone but further increased after efferocytosis of PMNs in PMs but was not affected by efferocytosis in BMDMs in response to *P. aeruginosa* (fig. S7F). Next, we tested human primary AMs (hAMs) taken from BALF of patients with uninflamed lungs. Like murine macrophages, hAMs failed to produce mtROS in response to bacteria after MPO pre-stimulation. Moreover, genipin prevented MPO-mediated impairment of mtROS generation, whereas α-KG restored MPO-mediated effects in the presence of genipin (Fig. 7D). Furthermore, human IL-1β secretion was reduced in response to bacteria after MPO pre-treatment, whereas human IL-6 was not affected (fig. S7G).

**Fig. 7.**
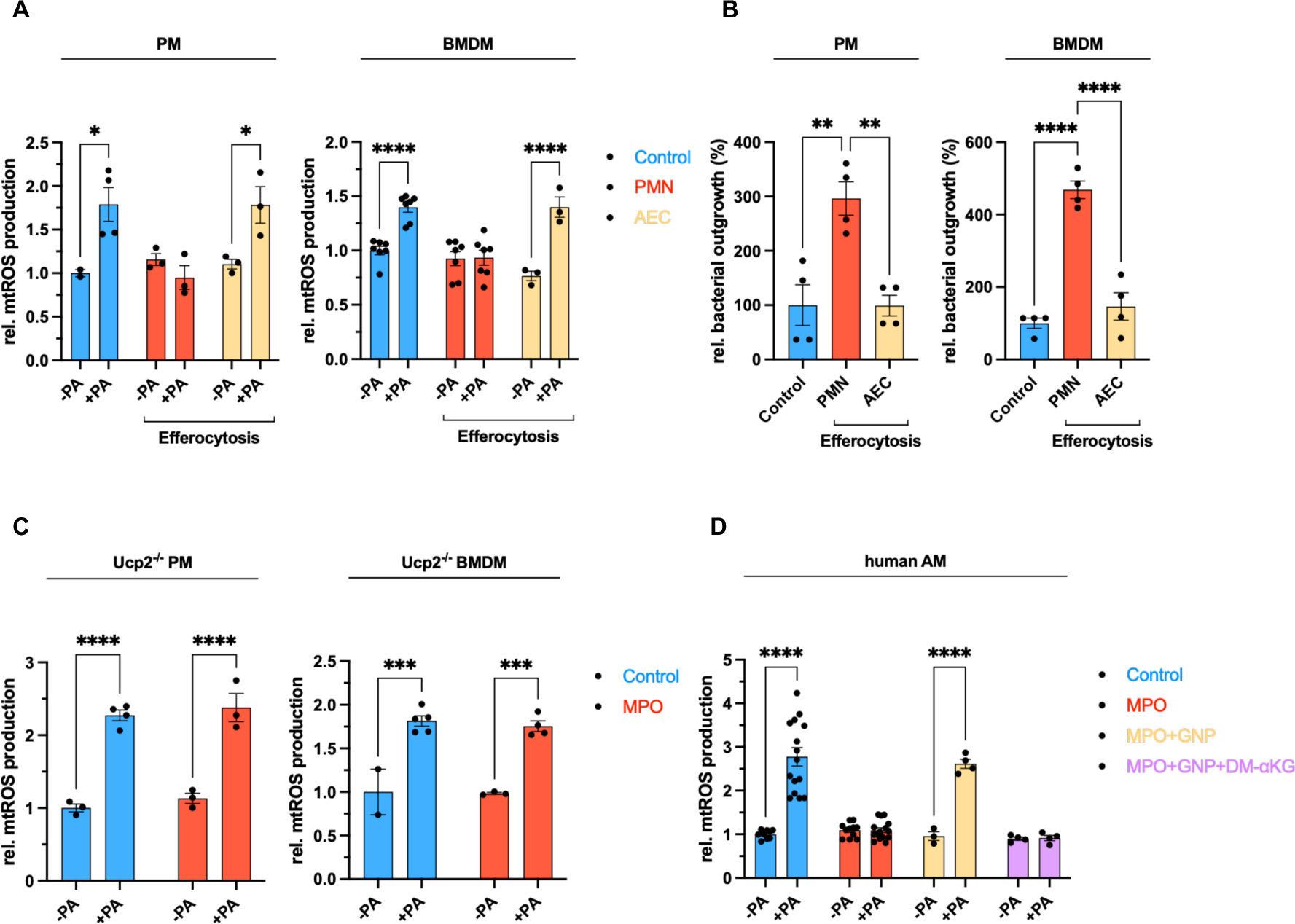
MPO impairs bactericidal properties of different macrophages across species via UCP2. (**A**) mtROS measurement in PMs (left) and BMDMs (right) after efferocytosis of apoptotic PMNs or AECs for 4 hours in response to PA for 6-8 hours (n=3-7 replicates/group, pooled from 3-5 mice/group and 2-3 independent experiments). (**B**) Bacterial killing capacity (*P. aeruginosa*) of PMs (left) and BMDMs (right) after efferocytosis of apoptotic PMNs or AECs for 4 hours in response to *P. aeruginosa* for 6-8 hours (n=4 replicates/group, pooled from three mice/group and two independent experiments). (**C**) mtROS measurement in AMs from WT and *Ucp2*^−/−^ PMs (left) and BMDMs (right) after pre-treatment with MPO 0.5μM for 3 hours in response to PA for 6-8 hours (n=3-5 replicates/group, pooled from 3-4 mice/group and two independent experiments). (**D**) mtROS measurement in human AMs after pre-treatment with MPO 0.5μM for 3 hours +/− GNP 100μM for 1 hour concomitantly +/− DM-α-KG (αKG) 1mM for 1 hour in response to PA (n=3-15 replicates/group, pooled from 4 patients and 4 independent experiments). *p < 0.05, **p < 0.01, ***p < 0.001, ****p < 0.0001 by one-way ANOVA with Dunnet’s multi comparisons test (**B**) or two-way ANOVA with Sidak’s multiple comparisons test (**A**, **C** and **D**). Data are shown as mean ± s.e.m.

Thus, MPO is a conserved signal in macrophages across species to blunt antibacterial responses via UCP2.

### MPO boosts efferocytosis of neutrophils in different macrophage subsets while limiting bacterial control *in vivo*

Efferocytosis promotes further uptake of apoptotic cells, referred to as “continued efferocytosis” (*37–39*). In line, we found that AMs exhibited an increased capacity to ingest apoptotic PMNs during resolution of inflammation (i.e., 24 hours after instillation of acid) (Fig. 8A). UCP2 was shown to decrease the MMP to enhance the continued clearance of apoptotic cells in macrophages (*12*); therefore, we tested the impact of cell-type specific efferocytosis in the context of an altered MMP due to bacterial stimulation. As expected, efferocytosis of both PMNs and AECs enhanced the uptake of apoptotic PMNs in a second round of efferocytosis in AMs (Fig. 8B, blue bars). Notably, stimulation with bacteria after the first round of efferocytosis with AECs and in control AMs (no efferocytosis) reduced ingestion of apoptotic PMNs in the second or first round, respectively (Fig. 8B, left and right red bar). In stark contrast, PMN efferocytosis caused a feed-forward loop of continued efferocytosis despite the presence of bacteria, as evidenced by a sustained efferocytosis level in the second round (Fig. 8B, middle red bar). Since MPO increases UCP2 expression, we speculated that MPO might foster continued efferocytosis. Indeed, MPO alone mimicked the effects of PMN efferocytosis in the presence of bacteria during the second round of ingestion of apoptotic cells (Fig. 8C). Importantly, *Ucp2*^−/−^ AMs did not enhance efferocytosis in response to MPO (Fig. 8D). Murine PMs and BMDMs (fig. S8A), as well as hAMs (Fig. 8E), similarly increased the ingestion of apoptotic PMNs after MPO treatment. To corroborate our findings *in vivo*, we performed acid aspiration followed by secondary bacterial infection with *P. aeruginosa* for 10 hours with or without i.t. pre-treatment with genipin (Fig. 8F). In line with our *in vitro* and *ex vivo* findings, pre-treatment with genipin increased the number of apoptotic cells per lung surface compared to control (Fig. 8G). To test the relevance of our findings on disease outcome in clinically relevant models, we performed second-hit models using acid aspiration or IAV infection as a first hit, followed by secondary bacterial infection with *P. aeruginosa* or *S. pneumoniae,* respectively as a second hit (Fig. 8H and Fig. 8I, respectively). Genipin pre-treatment restored bacterial control during secondary *P. aeruginosa* infection after acid aspiration (Fig. 5J) and improved survival in the same model (Fig. 8J). Similarly, genipin pre-treatment strongly reduced the bacterial load when mice were superinfected with *S. pneumoniae* during IAV pneumonia (Fig. 8K), consequently improving survival in this model (Fig. 8L).

**Fig. 8.**
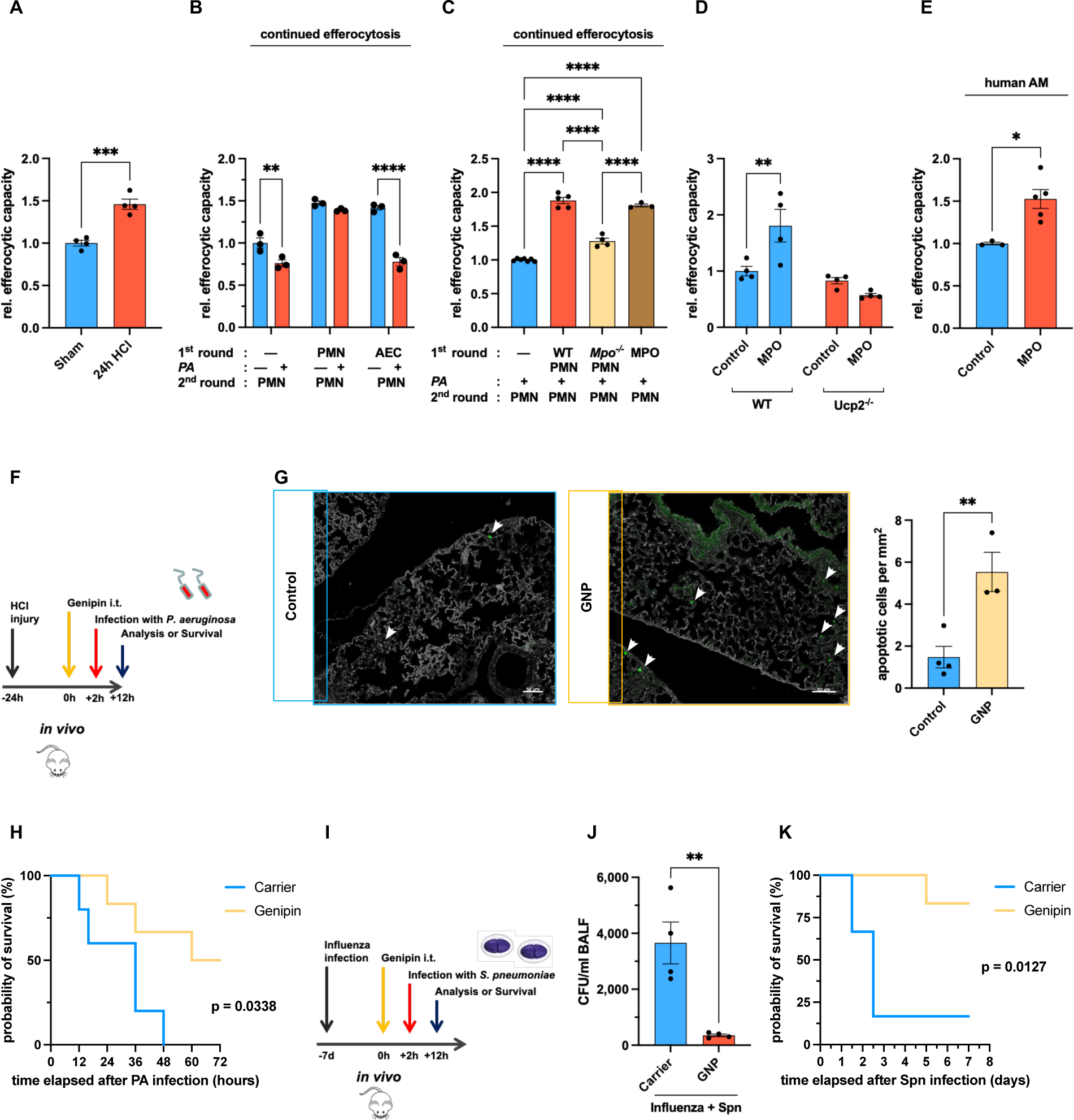
MPO prioritizes efferocytosis through UCP2 in different macrophage subsets across species while restricting bacterial control *in vivo*. (**A**) *Ex vivo* flow-cytometric analysis of AMs 24 hours after acid aspiration for their efferocytic capacity as determined by their ability to engulf apoptotic pre-stained PMNs within 1 hour compared to sham treatment (n=4 replicates/group, pooled from 4-6 mice/group and two independent experiments). (**B**) Continued efferocytosis *in vitro*: AMs were incubated with either apoptotic PMNs or AECs compared to control (no efferocytosis) for 4 hours +/− PA for 6-8 hours (1^st^ round = blue bars), AMs of all conditions were then incubated with apoptotic pre-stained PMNs for 1 hour to determine the capacity to engulf pre-labeled apoptotic PMNs (efferocytic capacity) by flow cytometry (2^nd^ round = red bars) (n=3 replicates/group, pooled from four mice/group and two independent experiments). (**C**) Continued efferocytosis: AMs were incubated with either apoptotic WT PMNs or *Mpo*^−/−^ PMNs compared to control (no efferocytosis) for 4 hours or pre-treated with MPO for 3 hours. All conditions were then stimulated with PA for 6-8 hours, followed by incubation with apoptotic pre-stained PMNs for 1 hour to determine the efferocytic capacity by flow cytometry (n=3-7 replicates/group, pooled from 4-8 mice/group and two independent experiments). (**D**) AMs of WT or *Ucp2*^−/−^ mice were treated with MPO 0.5µM for 3 hours compared to control. All samples were subsequently incubated with pre-stained apoptotic PMNs. Efferocytic capacity was then quantified by flow cytometry (n=4 replicates/group, pooled from five mice/group and two independent experiments). (**E**) hAMs were treated with human MPO 0.5µM for 3 hours compared to control before incubating all samples with pre-stained apoptotic human PMNs to determine efferocytic capacity (n=3-5 replicates/group, pooled analysis from 4 patients and four independent experiments). (**F**) Experimental scheme for (G and H). (**G**) Left: TUNEL-positive apoptotic cells (white arrowheads), depicted are representative images of (n=3-4 mice/group, pooled from two independent experiments); Right: Quantification of TUNEL-positive apoptotic cells. (**H**) Survival analysis after treatment as depicted in **f** (n=6 mice/group, pooled from two independent experiments). (**I**) Experimental scheme for (**J** and **K**). (**J**) Bacterial outgrowth in BALF 7 days after influenza infection followed by treatment with GNP 300µM intratracheally for 2 hours compared to control (DMSO) and subsequent bacterial superinfection with *S. pneumoniae* for 10 hours (n=4 mice/group, pooled from two independent experiments). (**K**) Survival analysis 7 days after influenza infection followed by treatment with GNP 300µM intratracheally for 2 hours compared to control (DMSO) and subsequent bacterial superinfection with *S. pneumoniae* for 10 hours (n=6 mice/group, pooled from two independent experiments). *p < 0.05, **p < 0.01, ***p < 0.001, ****p < 0.0001 by Student’s t-test (**A**, **E**, **G** and **H**, **J**) or one-way ANOVA with Dunnet’s multi comparisons test (**C**) or two-way ANOVA with Sidak’s multiple comparisons test (**B**, **D**). Survival distributions were followed up by a log-rank test (Mantel-Cox test) (**H**, **K**). Data are shown as mean ± s.e.m.

In conclusion, neutrophil-derived MPO facilitates the continued clearance of apoptotic cells in different macrophage subsets while restricting bacterial control *in vivo* mediated through a UCP2-controlled immunometabolic reprogramming (fig. S8B).

## Discussion

Macrophages are appreciated as orchestrators of inflammation by swiftly adopting functional phenotypes. However, mechanisms that regulate putatively antagonistic macrophage effector functions (e.g., pro-inflammatory/antibacterial versus anti-inflammatory/pro-resolving) are less clear. Here, starting from an unbiased *in vivo* approach, we uncovered that neutrophil efferocytosis reprograms mitochondrial metabolism to switch alveolar macrophages to a pro-resolution phenotype at the cost of bacterial control.

Mechanistically, UCP2 was reported to have mild uncoupling properties by increasing proton conductance of the inner mitochondrial membrane of the respiratory chain upon specific activation, diminishing mtROS release (*30, 40*). However, a mounting body of evidence highlights the role of UCP2 as a metabolic carrier (*33, 34, 41*). UCP2 was demonstrated to export C4 metabolites out of mitochondria, thereby limiting the oxidation of acetyl-CoA-producing substrates such as glucose while favoring glutaminolysis (*36*). By exporting C4 compounds from mitochondria, UCP2 reduces the redox pressure on the ETC and, hereby, mtROS release (*36*). Here, we show that enhanced UCP2 expression by MPO in macrophages precludes a hyperpolarized MMP and consecutive mtROS release in response to bacteria, which is reversed by inhibiting canonical glutaminolysis. Furthermore, a mitochondrial substrate oxidation stress test and UCP2-dependent increased glutamine consumption after MPO stimulation in response to bacteria demonstrated the importance of glutaminolysis in AMs. Therefore, we suggest that metabolic regulations by UCP2 are crucial in this context.

Increased UCP2 expression by MPO *in vitro* or during resolution of lung injury *ex vivo* was not paralleled by increased UCP2 transcripts, which is in agreement with previous observations (*42*). UCP2 has an unusual short half-live (*43*), and translational regulation by glutamate enables its fast upregulation (*44, 45*). Thus, it is tempting to speculate that the increased glutamine uptake triggered by bacteria after MPO stimulation promotes UCP2 expression. However, the mechanisms regulating UCP2 expression, degradation, or possible posttranslational modifications in general, and specifically by MPO remain to be explored in further studies.

Uptake of apoptotic cells blocks pro-inflammatory signaling pathways in macrophages (*46*). We here show that cell type-specific efferocytosis of neutrophils impacts secretion of IL-1β and partly IL-6 by a lack of mtROS in response to bacteria in AMs. In line, mtROS were previously reported to trigger IL-1β release via inflammasome activation (*47*) and to promote activation of mitogen-activated protein kinases (MAPKs) supporting IL-6 secretion (*29*). IL-6 levels were consistently diminished in settings of reduced mtROS release in AMs and BMDMs, but not in murine PMs and hAMs, manifesting cell-type and species-specific differences.

In contrast, IL-10 secretion by AMs was neither influenced by MPO nor by mtROS itself but was observed after PMN efferocytosis upon bacterial encounter. The mechanism of enhanced, anti-inflammatory IL-10 release in this setting remains to be investigated. However, this effect is consistent with the overall notion that PMN efferocytosis primes for resolution of inflammation, while limiting pro-inflammatory responses in the presence of bacteria. Overall, we found a conserved effect of PMN efferocytosis on immune functions, causing a distinct shift from pro- to anti-inflammatory cytokines.

Besides, intentionally decreased release of mtROS in the center of neutrophils where phlogistic cargo accrues might have the purpose to further limit organ damage. Along these lines, scavenging of mtROS had protective effects in LPS-endotoxemia (*48*) and influenza pneumonia (*49*) by limiting the release of pro-inflammatory cytokines, and reducing apoptosis and necrosis of neutrophils.

Improved efferocytosis after ingestion of apoptotic cells, a phenomenon termed *continued efferocytosis,* is a well-established process (*50*). Accordingly, ingestion of apoptotic PMNs or AECs enhanced continued efferocytosis *in vitro*. However, AMs that had ingested AECs exhibited a “back to baseline” efferocytosis upon bacterial challenge, while PMN efferocytosis maintained a high efferocytic capacity of AMs even in the presence of bacteria in an MPO-dependent manner. Our data not only support the overall concept that PMN efferocytosis initiates and drives resolution of inflammation (*51–53*), but furthermore reveal that ingestion of apoptotic PMNs prioritizes continued efferocytosis over antibacterial responses via UCP2 in macrophages. This functional fate decision makes sense, given that accrual of necrotic PMNs can promote inflammation and tissue damage (*54, 55*). Intriguingly, MPO in PMNs was found to protect mice during sterile endotoxemia (*56*), and MPO deficiency increased atherosclerotic lesions (*57*). In both studies the mechanism remained unsolved. It is tempting to speculate that a functional switch to a pro-resolution/anti-inflammatory state induced by MPO might play a role. In yet another interesting study, Faas et al. recently discovered that UCP2 promotes repair mechanisms in macrophages in response to the alarmin IL-33 in a model of muscle injury (*58*). Thus, different danger signals (i.e., MPO, IL-33) increase UCP2 expression to improve resolution properties of macrophages.

In summary, we delineated a conserved mechanism of MPO-dependent mitochondrial reprogramming that restricts the functional plasticity of macrophages to prioritize tissue-protective properties. These findings were conserved across species, macrophage subsets, and bacterial pathogens. While this is advantageous in most circumstances, it creates a window of opportunity for colonizing bacteria during the resolution of inflammation, which might result in severe gram-negative pneumonia after aspiration or gram-positive pneumonia after viral infection. Considering an increasing number of clinical studies that evaluate the therapeutic potential of cell therapies using apoptotic cells (*59–62*), our results stress the importance to discern the effects of different apoptotic cell preparations on macrophages. Furthermore, our data suggest that modulation of canonical glutaminolysis or UCP2 might constitute a therapeutic approach to instruct macrophages for improved host defense or repair functions after aspiration or during viral pneumonia.

## List of abbreviations

AEC: Alveolar epithelial cell
α-KG: Alpha-ketoglutarate
AM: Alveolar macrophage
BALF: Broncho-alveolar lavage fluid
BMDM: Bone-marrow-derived macrophage
cROS: Cytosolic reactive oxygen species
ECAR: Extracellular acidification rate
EGCG: Epigallocatechin-3-gallate
GNP: Genipin
HCl: Hydrochloric acid
i.n.: intranasally
i.t.: intratracheally
IL: Interleukin
KEGG: Kyoto Encyclopedia of Genes and Genomes
*K. pneumoniae/KP*: *Klebsiella pneumoniae*
MDM: Recruited inflammatory Monocyte-derived macrophage
MitoT: MitoTempo
mtROS: Mitochondrial reactive oxygen species
MPO: Myeloperoxidase
OCR: Oxygen consumption rate
OXPHOS: Oxidative phosphorylation
*P. aeruginosa/PA*: *Pseudomonas aeruginosa*
PM: Peritoneal macrophage
PMN: Polymorphonuclear leukocyte
ROS: Reactive oxygen species
*S. pneumoniae/SP*: *Streptococcus pneumoniae*
TNF-α: Tumor necrosis factor-α
WT: Wildtype

## Acknowledgments

We thank Larissa Hamann, Stefanie Jarmer, Florian Lück, and Julia Stark for their excellent technical assistance. We are grateful to Sylvia Knapp for critically reading the manuscript and Elena Pohl for fruitful discussions.

## Funding

German Research Foundation grant “KFO309”, project number 284237345 (SH)

German Research Foundation grant “SFB-TR84”, project number 114933180 (SH)

LOEWE grant “diffusible signals” (UM and SH)

German Center for Lung Research (DZL) (UM and SH)

Cardiopulmonary Institute (CPI) - DFG EXC 2026 (SH)

## Author contributions

Conceptualization: JB, UM, SH

Methodology: JB, MW, UM, SH

Investigation: JB, MW, ME, CM, MRF, ML, IK, IA, SM

Bioinformatics and Statistics: JB, JW, UM

Visualization: JB, UM, JW, IA

Funding acquisition: UM and SH

Project administration: UM and SH

Supervision: UM and SH

Writing – original draft: JB and UM

Writing – review & editing: JB, UM, SH, IV-A, LK, RS, NS

## Supplementary Material

### Summary

Materials and Methods

Resources Table

References (63–67)

Figure S1. Functional and transcriptional properties of AMs during acid aspiration.

Figure S2. Determining mitochondrial reactive oxygen species production by AMs after acid aspiration.

Figure S3. Cell-type specific efferocytosis of PMNs precludes mtROS generation in response to bacteria in AMs.

Figure S4. PMN-derived MPO mediates alterations of mitochondrial signaling in AMs.

Figure S5. MPO-mediated increase of UCP2 precludes mtROS release in response to bacteria.

Figure S6. MPO drives canonical glutaminolysis through UCP2.

Figure S7. MPO acts via UCP2 to impair bacterial control in different macrophages across species.

Figure S8. MPO acts via UCP2 to improve efferocytic capacity in different macrophages.

## Materials and Methods

### Study design

The study was designed to identify local environmental cues that determine the transcriptome, immunometabolic phenotype, and immune effector function of AMs during resolution of inflammation influencing the outcome during subsequent secondary bacterial infection. Subjects or other experimental units were assigned randomly except that 10- to 12-week-old, age- and sex-matched mice were used for all experiments unless otherwise specified. Littermates were included where possible. The experiments and assessment of outcomes were performed in an unblinded fashion. Descriptions of experimental replicates are found in figure legends and/or subsections in Materials and Methods.

### Animals

Female C57BL/6N wildtype mice were purchased from Charles River Laboratories at the age of 8-12 weeks. *Ucp2*^−/−^ mice (B6.129S4-Ucp2^tm1Lowl^) were kindly provided by the laboratory of Natascha Sommer (University of Giessen, Hesse, Germany). Breeding pairs of *Mpo*^−/−^ mice (B6.129X1-Mpo^tm1Lus/J^) had been purchased from Charles River. All mice were housed in a specific pathogen-free environment. Animal experiments were conducted according to the legal regulations of the German Animal Welfare Act (Tierschutzgesetz) and approved by the regional authorities of the State of Hesse (Regierungspräsidium Giessen) and the Institutional Commission for the Care and Use of Laboratory Animals (CICUAL), Buenos Aires, Argentina.

### Induction of acid aspiration

Mice were anesthetized with isoflurane (Sigma-Aldrich, Germany). Acid pneumonitis was induced by instillation of 50µl of 0.1M HCl intratracheally (i.t.) (Sigma-Aldrich, Germany). Sham-treated controls received 50µl of 0.9% NaCl (Braun, Germany) intratracheally (i.t.).

### Bacterial superinfection of Influenza A virus pneumonia with S. pneumoniae

*S. pneumoniae* serotype 3 was cultured in Todd-Hewitt Broth (THY) containing 10% heat-inactivated FCS at 37°C mid-log phase. Inoculum was determined by serial dilutions on blood agar plates. Mice were anesthetized with isoflurane (Sigma-Aldrich, Germany). Then, 100ffu of Influenza A virus (PR8) diluted in 70μl sterile PBS−/− were instilled i.t.. Seven days after Influenza A virus infection, mice were treated with 50μl of 300μM Genipin or DMSO (control for Genipin) i.t. and two hours later intranasally infected with 30 colony forming units (CFU) Streptococcus pneumoniae (diluted in 50μl NaCl 0,9%).

### Bacterial pneumonia with *Pseudomonas aeruginosa*

*Pseudomonas aeruginosa (P. aeruginosa)*, strain PA103, was cultured in Luria broth (LB) medium (Roth) at 37°C with aeration and harvested at the mid-log phase. Inoculum was determined by serial dilutions on blood agar plates, and mice were infected with 2-3×10^4^ colony forming units (CFU) of *P. aeruginosa* instilled in 50µl NaCl 0,9% intranasally at indicated time points. Before inoculation, mice were short-term anesthetized by inhalation of isoflurane.

### Primary cell harvest and culture

All murine alveolar Macrophages (AMs) were obtained from BALF. Mice were sacrificed, the trachea was exposed, incised, and intubated with a blunt 21G cannula. BAL was performed by successively instilling seven times 1mL of cold PBS−/− plus EDTA intratracheally. Recovery volume was centrifuged at 500g for 10 minutes. The supernatant was discarded while the cell pellet was resuspended in an RPMI-based medium containing 2.5% HEPES, 2% FCS, and 1% Penicillin/Streptomycin/L-Glutamine, and cultured at 37°C under 5% CO_2_. After cells had adhered, non-adherent cells were washed away.

Human BALF with a macrophage purity > 90% obtained from patients for diagnostic purposes was applied to isolate human alveolar macrophages (hAMs). BAL samples used were from non-smoking patients without chronic pulmonary diseases who provided written consent to the use of biomaterial using the consent form of the DZL (Deutsches Zentrum für Lungenforschung). The project was reviewed and is covered under the University of Giessen ethics committee decision (AZ 58/15) and was performed according to the appropriate regulations and the Declaration of Helsinki. BALF was centrifuged before cells were resuspended in an RPMI-based medium containing 2.5% HEPES, 2% FCS, and 1% Penicillin/Streptomycin/L-Glutamine, and cultured at 37°C under 5% CO_2_. After cells had adhered, non-adherent cells were washed away.

Murine peritoneal macrophages (PMs) were taken from peritoneal lavage fluid. Mice were euthanized, and the inner skin lining of the peritoneal cavity was exposed. The peritoneal lavage was performed by instilling 7-10mL of HBSS through a 27G needle intraperitoneally. The peritoneum was gently massaged before collecting the peritoneal fluid with a 23G needle with the mouse placed in a lateral decubitus position. Recovery volume was centrifuged, and the supernatant was discarded while PMs were purified according to manufacturer’s instruction using a magnet purification kit by selecting F4/80^+^ cells (Miltenyi Biotec; Macrophage Isolation Kit, Mouse, Peritoneum). Finally, purified PMs were resuspended in an RPMI-based medium containing 10% FCS and 1% Penicillin/Streptomycin/L-Glutamine and cultured at 37°C under 5% CO_2_.

Murine bone marrow-derived macrophages (BMDMs) were generated from femoral and tibial bone marrow. After mice had been sacrificed, femoral and tibial bones were isolated by removing attached muscles and tissue. The collected bone marrow was further passed through a 40 µm cell strainer to remove gross particles before centrifuging. The supernatant was discarded while the cell pellet was resuspended in an RPMI-based medium containing 10% FCS, 30ng/mL M-CSF, and 1% Penicillin/Streptomycin/L-Glutamine, and cultured at 37°C under 5% CO_2_. After cells had adhered, non-adherent cells were washed away, and cells were grown for 7d by adding fresh medium on days 3 and 5.

Murine bone marrow-derived neutrophils (PMNs) were isolated from femoral and tibial bone marrow as described above. Subsequently, cells were spun down at 500g for 10 minutes and resuspended in MACS buffer to purify PMNs with a magnetic purification kit according to manufacturer’s instruction (Miltenyi Biotec; Neutrophil Isolation Kit, Mouse).

### In vitro cell culture

Murine lung alveolar epithelial cells (AECs), MLE-12 (ATCC: CRL-2110), were grown in a T-75 cell culture flask supplemented with a DMEM-based medium containing 10% FCS, ITS (Insulin 0.005 mg/ml, Transferrin 0.01 mg/ml, Sodium selenite 30 nM), 1% Penicillin/Streptomycin/L-Glutamine at 37°C under 5% CO2.

### Ucp2 transcription

RNA was isolated using RNeasy Micro Kit (QIAGEN) according to manufacturer’s protocol. 250ng of isolated RNA was reverse-transcribed into cDNA. Quantitative PCR was performed with SYBR green I (Invitrogen) in the AB StepOnePlus Detection System (Applied Bioscience). mRNA amounts are presented as fold change normalized to b-actin expression as well as the untreated or sham control. The following primers were used: murine β-actin (forward primer, 5′-ATGGGAAGCCGAACATACTG-3′; reverse primer, 5′-CAGTCTCAGTGGGGGTGAAT-3′) and murine Ucp2 (forward primer, 5′-AAGGGCTCAGAGCATGCAG-3′; reverse primer, 5′-TGGAAGCGGACCTTTACCAC-3′).

### Efferocytosis

Murine bone marrow-derived PMNs were resuspended at 5-10×10^5^ cells/ml in an RPMI-based medium containing 10% FCS and 1% Penicillin/Streptomycin/L-Glutamine before 0.2 µM staurosporine was added. The cell suspension was incubated for 6 hours at 37°C under 5% CO_2_. MLE-12 cells were treated with 0.4µM staurosporine for 24 hours at 37°C and 5% CO_2_ in MLE cell culture medium. A subsequently performed flow cytometry-based annexin V/7-AAD staining confirmed equal amounts of apoptotic cells (75 +/− 5%) and less than 10% dead cells for both PMNs and MLE-12 cells. Finally, both cell types were centrifuged and washed twice to remove extracellular staurosporine.

For all efferocytosis experiments, three apoptotic cells (PMNs or AECs) per macrophage were added, meaning an apoptotic cell to efferocyte ratio of 3:1, and incubated for 4 hours before apoptotic cells were removed. These efferocytosis experiments refer to efferocytosis as a macrophage treatment.

For determining the efferocytic capacity of primary macrophages, i.e., their capacity to clear apoptotic cells, apoptotic PMNs were pre-incubated with 5µM CellTrace Calcein Red-Orange for 30 min at a cell concentration of 1-10×10e6 cells/mL. After washing, three apoptotic cells per macrophage were added, meaning an apoptotic cell to efferocyte ratio of 3:1, incubated for 60 minutes before apoptotic cells were removed. Macrophages were then subjected to flow cytometric analysis to quantify the fluorescence signal of CellTrace Calcein Red-Orange in macrophages. The mean fluorescence intensity correlates with the capacity of macrophages to clear apoptotic bodies (PMNs). The mean intensity signal of the control treatment was used to normalize the efferocytic capacity of each individual experiment.

Quantification of continued efferocytosis, referring to the continued clearance of apoptotic cells, was performed in a two-step model. During the first round of efferocytosis, macrophages were incubated with either apoptotic PMNs or AECs for 4 hours compared to control (no efferocytosis). In a second round of efferocytosis, macrophages of all conditions were then incubated with apoptotic pre-stained PMNs, as described above, to determine their efferocytic capacity.

### Bacterial killing assay

Primary macrophages were cultured for 3h before analysis. Afterward, each well was washed four times with NaCl 0,9% (Braun, Germany) to remove non-adherent cells, and RPMI medium supplemented with 2% FCS and 2.5% HEPES buffer was added afterward. *P. aeruginosa* (PA103) and *S. pneumoniae* (S23) were grown as indicated. *K. pneumoniae* (ATCC 700721) was grown at mid-log phase in LB medium. After thorough washing, bacteria were added with a multiplicity of infection (MOI) of approximately 5 (*P. aeruginosa*), 10 (*K. pneumoniae*), and 250 (*S. pneumoniae*), and both plates were incubated for 1h (*P. aeruginosa* and *K. pneumoniae*) or 10 min (S. pneumoniae) with antibiotic-free medium. Then both plates were washed four times again. One plate was immediately lysed with distilled water (Braun, Germany) to assess bacterial uptake. The other plate was left for an additional 1.5h (*P. aeruginosa*), 3h (*K. pneumoniae*), or 20 min (*S. pneumoniae*) and subsequently lysed to assess bacterial killing (relative to bacterial uptake). Serial dilutions of the lysate were incubated overnight on blood agar plates to quantify viable bacteria in each condition.

### Assessment of mtROS, cROS, and MMP

Primary macrophages were seeded with 1.5*10^5^ live cells per well in a 48-well plate (Greiner BIO-ONE, Germany). After adherence, each well was washed once with PBS, and heat-killed PA103, *K. pneumoniae,* or *S. pneumoniae* (MOI 100) was added for 6-8 hours. Subsequently, wells were washed twice, and cells were incubated with 3.5µM MitoSOX Red (ThermoFisher, United States) for 20 minutes for the assessment of mtROS, 10µM CM-H2DCFDA (Thermo Fisher Scientific) for 15 minutes for the assessment of cROS, or 7.7µM JC-1 (ThermoFisher) for 30 minutes for the assessment of the MMP. After washing, cells were immediately transferred to FACS tubes (pluriSelect, Germany) and stored on ice. The analysis was done with a BD LSRFortessa and FACS Diva Software (Becton Dickinson, United States). Data were analyzed with FlowJo Version 10.6.2.

### Cytokine Measurements

Primary macrophages were cultured for 3h prior to analysis and incubated with heat-killed *P. aeruginosa* (MOI 100) for 6-8 hours. Protein concentrations of selected cytokines and chemokines were measured in supernatants of macrophages using a BioPlex MAGPIX Multiplex Reader (BIO-RAD, United States) according to manufacturer’s instructions. Data were analyzed with Bio-Plex Data Pro software.

### Flow cytometry and cell sorting

BAL was centrifuged and subsequently counted with an NC-250 NucleoCounter (ChemoMetec, Denmark). Cells were resuspended in blocking reagent (Gamunex, Grifolis, Spain) and indicated antibodies (listed in the Key Resources Table) for 30 min 4°C, cell suspensions were centrifuged again and resuspended in FACS buffer containing phosphate buffered saline, supplemented with 10% FCS, 0.1% sodium azide. Sytox live-dead staining was added immediately before analyzing a sample. For annexin V staining, cells were resuspended in annexin V binding buffer (10 mM HEPES, 140 mM NaCl, and 2.5 mM CaCl2) at a concentration of 0.25-1.0×10e7 cells/mL. Then, 100µL of the cell suspension were transferred to a 5mL FACS tube. Subsequently, 5µL of each Pacific Blue Annexin V and 7-AAD was added to the staining solution. The solution was vortexed gently and incubated for 15 min at room temperature in the dark. Finally, 400µl of Annexin V binding buffer was added. Flow cytometric analysis was done with BD LSRFortessa and BD FACSAria III cell sorter using FACS Diva Software (Becton Dickinson, United States). Data were analyzed with FlowJo Version 10.6.2. Live/dead staining was performed with Sytox Blue Pacific Blue (BioLegend) or 7-AAD.

### Bulk mRNA sequencing

Purified total RNA was amplified using the Ovation PicoSL WTA System V2 kit (NuGEN Technologies, Bemmel, Netherlands). Per sample, 2µg amplified cDNA was Cy-labeled using the SureTag DNA labeling kit (Agilent, Waldbronn, Germany). The Cy5-labeled cDNA was hybridized overnight to 8 x 60K 60mer oligonucleotide spotted microarray slides (Agilent Technologies, design ID 074809). Hybridization and subsequent washing and drying of the slides was performed following the Agilent hybridization protocol. The dried slides were scanned at 2 µm/pixel resolution using the InnoScan is900 (Innopsys, Carbonne, France). Image analysis was performed with Mapix 8.2.5 software, and calculated values for all spots were saved as GenePix results files. Stored data were evaluated using the R software (Team, 2007) (3.5.1) and the limma package (*63*) (3.30.13) from BioConductor (*64*). Gene annotation was supplemented by NCBI gene IDs via bioMart (*65*) (accessed 2018-03-08). Mean spot signals were background corrected with an offset of 1 using the NormExp procedure on the negative control spots. The logarithms of the background-corrected values were quantile-normalized (*66*). The normalized values were then averaged for replicate spots per array. From different probes addressing the same NCBI gene ID, the probe showing the maximum average signal intensity over the samples was used in subsequent analyses. Genes were ranked for differential expression using a moderated t-statistic. Pathway analyses were done using gene set tests on the ranks of the t-values. Gene sets were defined according to the KEGG database (*67*) (accessed 2018-03-08).

### Extracellular flux analyses

Real-time bioenergetic profiles of macrophages were observed by measuring oxygen consumption rate (OCR) using either Agilent Seahorse XF HS Mini or XFe96 analyzer (Seahorse Bioscience, Agilent Technologies, North Billerica, MA). Macrophages were plated on an 8-well XF HS Mini PDL and a 96-well XFe96 cell culture microplate at a density of 25,000 cells or 70,000 cells per well, respectively. After indicated treatments, extracellular flux analysis was performed as per manufacturer’s instructions.

In brief, cellular bioenergetics were investigated by applying an XF Mito Stress Test necessitating the sequential injection of 1.5µM oligomycin, 2.5µM carbonyl cyanide 4-(trifluoromethoxy) phenylhydrazone (FCCP) and 0.5/0.5µM rotenone/antimycin A.

To reveal the metabolic phenotype and function of macrophages, XF Substrate Oxidation Stress Test was conducted, combining pathway-specific inhibition of the primary mitochondrial substrates with a subsequent XF Mito Stress Test. Therefore, 2µM UK5099, 3µM BPTES, 4µM Etomoxir, and medium only (control) were injected concomitantly in Port A of separate wells to interrogate the contribution of glycolysis, glutaminolysis, or fatty-acid oxidation to mitochondrial respiration. Each of these injections were followed by an XF Mito Stress Test as described above. The relative importance of each of the mentioned pathways for feeding mitochondrial respiration was determined by the potential of the pathway-specific inhibitors to reduce maximal mitochondrial respiration in comparison to the uninhibited control meaning the greater the reduction of maximal respiration, the greater the cell relies on the specific pathway for fueling mitochondrial respiration and vice versa.

Data were analyzed with Seahorse Analytics (Agilent)

### Western Blot

Macrophages were treated as indicated, detached by adding Trypsin/EDTA for 10min at 37°C/5%CO2. Cells had been centrifuged at 500g for 10min before supernatant was discarded. Subsequently, cell pellet was resuspended in lysis buffer containing 900µL NP-40 buffer, 100µL Proteinase-Inhibitor Cocktail (PIC), and 3.4µL DDT. Samples were then incubated on ice for 30min while being puls-vortexed every 5-10min. Finally, cell debris was precipitated by centrifugation at maximum speed (>= 16,000g) for 15min at 4°C. Protein concentration was quantified by Bradford Assay according to manufacturer’s instructions. A sample volume containing 20-30µg of protein was mixed with loading buffer (900µL 4x Laemmli buffer + 100µL 2-Mercaptoethanol) in a ratio of 3:1. Samples were further boiled at 95°C for 5min before being cooled on ice again. 20-40µg of total protein per well were loaded on a gel (5-15%). The membrane was blocked at RT for 1h in 2% BSA in TBS-T 0.01%. Primary antibody solution was prepared by diluting UCP2 (D105V) Rabbit mAb 1:1000 in 2% BSA in TBS-T 0.01% and added to membrane overnight at 4°C. The membrane was washed three times with TBS-T before the secondary antibody (HRP anti-Rabbit 1:1000) was added for 1h at RT. Finally, imaging was performed. To detect the β-Actin loading control, the membrane was stripped for 30min at RT and subsequently incubated with primary and secondary antibody.

### TUNEL assay

In situ nick-end labeling of nuclear DNA fragmentation was performed with a TUNEL apoptosis detection kit (DeadEnd Fluorometric TUNEL system, Promega) according to the supplier’s instructions.

For the acquisition of fluorescence images of the TUNEL staining, we used the EVOS widefield microscope and the 10x air objective. Two channels were acquired per field of view. One corresponding to the emission of the TUNEL staining by setting the filter to the GFP position, and one using the transmitted light to capture the tissue brightfield image. Multiple tiles (fields of view) were captured and then stitched with the acquisition software of the EVOS system. Similarly, we captured the brightfield images of the H&E staining with the same EVOS microscope and used the 10x air objective again, using transmitted light and setting the camera to color mode. The whole lung slices were captured using multiple fields of view and stitching those with the acquisition software of EVOS.

The number of TUNEL-positive cells was counted from the widefield fluorescence images using a custom-made Fiji macro. The raw images were thresholded, and a binary mask was generated. Using the brightfield image, the boundaries of the lung section were delineated. With the “Analyse Particles” plugin, we counted the number of TUNEL-positive cells within the tissue selection. Their number was then normalized by the total tissue area and reported as the number of cells per mm2 tissue.

### Mitochondrial content *in vivo*

DNA from cell lysates of alveolar macrophages, harvested 24 hours after acid aspiration compared to sham treatment, was isolated using DNeasy Blood & Tissue Kit (QIAGEN) according to manufacturer’s protocol and normalized using NanoDrop (ThermoFischer Scientific). Quantitative PCR was performed with SYBR green I (Invitrogen) in the AB StepOnePlus Detection System (Applied Bioscience). Mitochondrial content was quantified by the ratio of mitochondrial DNA to genomic DNA. The following primers were used: genomic DNA (murine β2 microglobulin) (forward primer, 5′-ATGGGAAGCCGAACATACTG-3′; reverse primer, 5′-CAGTCTCAGTGGGGGTGAAT-3′) and mitochondrial DNA (forward primer, 5′-CTAGAAACCCCGAAACCAAA-3′; reverse primer, 5′-CCAGCTATCACCAAGCTCGT-3′).

### Mitochondrial content *in vitro*

Successful mitochondrial staining was confirmed by colocalization of anti-TOMM20 and MitoTracker deep red staining. TOMM20 is a translocase of the outer mitochondrial membrane. MitoTracker deep red contains a mildly thiol-reactive chloromethyl moiety and accumulates in mitochondria, but in addition, staining intensity might be influenced by the mitochondrial membrane potential. Therefore, mitochondrial content was finally quantified by the mean intensity of the anti-TOMM20 signal.

AMs were seeded equally on precision coverslips (diameter: 10mm; thickness: #1.5H) and subsequently treated with 0.5µm MPO for 2.5-3h. After that, mitochondria were stained with MitoTracker deep red 100nm for 30min at 37°C/5%CO2. Cells were washed three times with PBS 1X. Cell fixation was achieved by adding 4% PFA (pH 7.4) for 10min at 37°C. Cells were then washed three times with PBS 1X. 0.1% Triton X-100 in PBS 1X was applied to permeabilize cells at RT for 15min before rinsing three times with PBS 1X. Blocking was performed by adding 5% goat serum containing 0.1% sodium azide at RT for 60min, followed by staining with anti-TOM20 recombinant rabbit monoclonal antibody at a 1:1000 dilution in 5% goat serum containing 0.1% sodium azide overnight at 4°C. The next day, cells were washed three times with PBS 1X. Goat anti-rabbit recombinant secondary antibody Alexa Fluor 555 was diluted 1:1000 in 5% goat serum containing 0.1% sodium azide and added to the cells for 1h at RT protected from light. Cells were washed three times with 1X PBS-T. Nuclei were stained with DAPI 10μg/mL in PBS for 10 minutes and rinsed thrice with PBS-T 1X. Coverslips were finally air-dried as well as mounted with ProLong glass antifade mountant.

Confocal images were acquired with an SP8 confocal microscope (Leica Microsystems) equipped with a white light laser (WLL) and multiple Hybrid Detectors (HyD) using the LAS X software (v3.5.7). Stained samples were scanned sequentially using a PL APO 63xOil /1.40 NA objective with a voxel size of 28×28×200nm. The pinhole was kept closed to 0.7AU, and we applied a 2x live average to eliminate the noise. For the excitation of DAPI, we used 0.1% of a 405nm laser. For the excitation of TOM20 and MitoTracker deep red, we used 4.3% of a 553nm laser line or 6.7% of a 641nm laser line, respectively, derived from a white light laser set at 50% of its nominal power. For the detection of the emitted wavelengths, we used three Hybrid detectors set at 413-550nm, 560-636nm, and 650-780nm. To measure the mean intensity of TOMM20, we acquired confocal images with a lower magnification objective (20xAir/0.75NA) to capture as many cells as possible. All acquired z-stacks were subsequently cleared computationally using the Lightning module of the LAS X software.

The mean intensity of the TOMM20 marker as a surrogate marker for mitochondrial content was quantified from low magnification confocal images using ImageJ/Fiji as follows: Initially, we performed a maximum intensity projection for the acquired image stacks. The signal of the MitoTracker deep red was used to generate masks of cells upon filtering the generated projections with the median filter (radius 1.5) and thresholding using the Huang algorithm. The individual cells per field of view were marked with regions of interest (ROIs) using the “Analyse Particles” plugin of Fiji. Subsequently, the background pixel intensities of the TOM20 fluorescence channel were removed with the default thresholding algorithm, and for each cell-ROI, the mean fluorescence intensity was measured.

### Bioluminescent metabolic measurement

AMs were isolated from WT or Ucp2^−/−^ mice and seeded at a density of 5×10^4^ cells per well supplemented with 100μL cell culture medium. In the following, cells were treated as indicated. Supernatant of each well was taken up, diluted 50-fold and split into three while cells were lysed in 100μL lysis solution (50μL 0.6M HCl and 50μL 1M TRIS) before being split into three. From different fractions of supernatant and cell lysate concentration of glucose, lactate, and glutamine or glutamine and glutamate were quantified, respectively. Measurement was performed by applying Glucose-Glo (Glucose), Lactate-Glo (Lactate), and Glutamine/Glutamate-Glo (Glutamine and Glutamate) kits from Promega according to manufacturer’s instructions. For Glucose-Glo kit supernatant was further diluted three-fold. relative light units were measured by a luminometer (Flx800, Biotek). Actual concentrations were then quantified from standard curves of each metabolite.

### Statistical analysis

Statistical analyses were performed in GraphPad Prism v.9 software. Data are presented as mean +/− standard error of the mean (s.e.m.). p-values less than 0.05 were considered significant, whereas p-values more than 0.05 were considered non-significant and not indicated unless stated otherwise. The mean between two groups was analyzed using the two-sided Student’s t-test. Statistically significant differences between the means of three or more unrelated and independent groups were compared using the one-way ANOVA test. p-values were corrected by Dunnet’s multiple comparisons test when comparing different groups to one control group or Tukey’s multiple comparisons test when comparing all different groups with each other. The two-way and three-way ANOVA approaches were used to estimate how the mean of a quantitative variable changes according to the levels of two or three categorical variables, respectively. Šídák’s multiple comparisons test was further applied to adjust p-values. Survival distributions of two samples were followed up by a log-rank test (Mantel-Cox test). Data distribution was assumed to be normal, but this was not tested formally. Plated cells were allocated randomly to each treatment group; C57Bl/6 mice were assigned randomly to each treatment group. Data collection and analysis were not performed blind to the conditions of the experiments.

## Resources Table

### Machines and devices

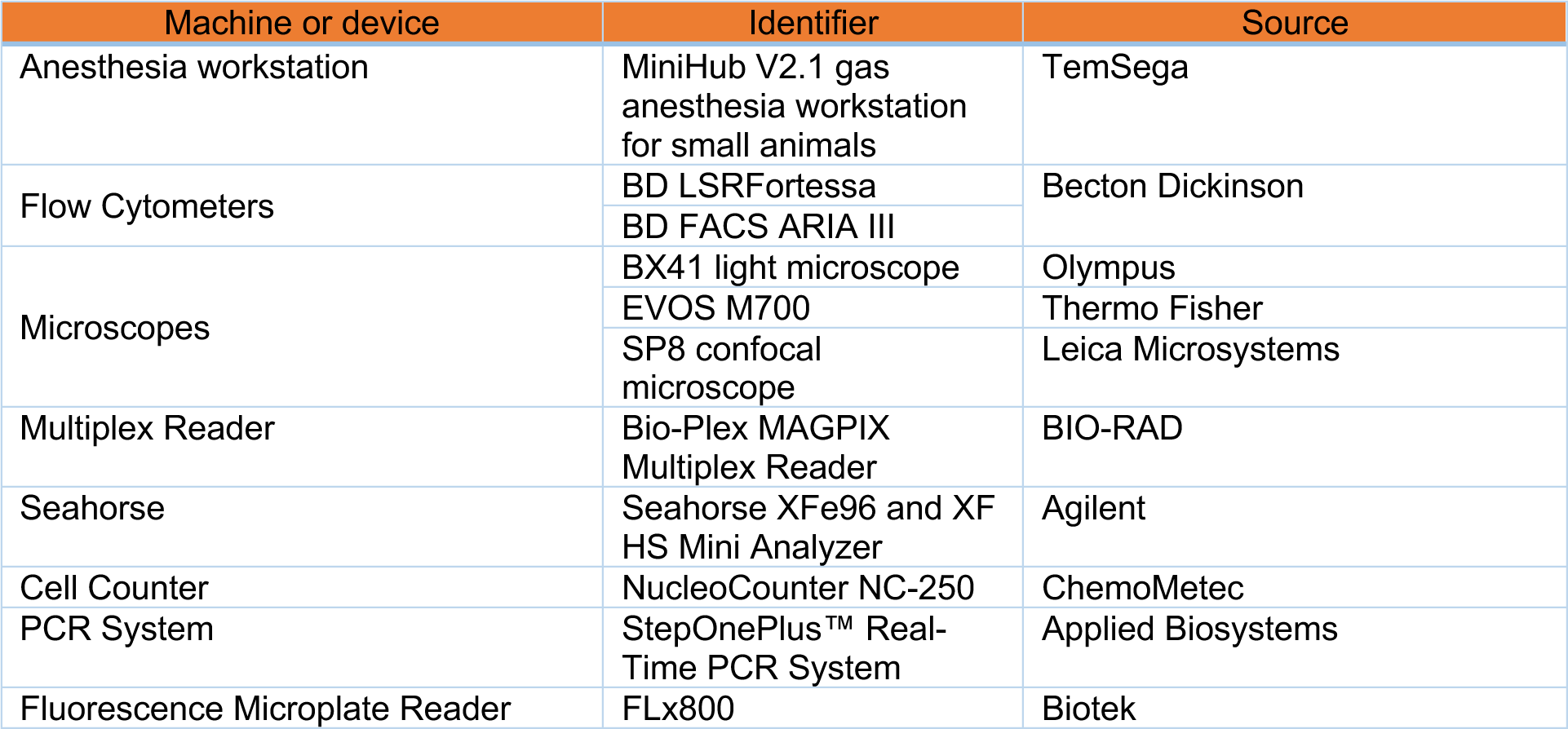

| Machine or device | Identifier | Source |
| --- | --- | --- |
| Anesthesia workstation | MiniHub V2.1 gas anesthesia workstation for small animals | TemSega |
| Flow Cytometers | BD LSRFortessa | Becton Dickinson |
|  | BD FACS ARIA III |  |
| Microscopes | BX41 light microscope | Olympus |
|  | EVOS M700 | Thermo Fisher |
|  | SP8 confocal microscope | Leica Microsystems |
| Multiplex Reader | Bio-Plex MAGPIX Multiplex Reader | BIO-RAD |
| Seahorse | Seahorse XFe96 and XF HS Mini Analyzer | Agilent |
| Cell Counter | NucleoCounter NC-250 | ChemoMetec |
| PCR System | StepOnePlus™ Real-Time PCR System | Applied Biosystems |
| Fluorescence Microplate Reader | FLx800 | Biotek |

### Mitochondrial probes

| Mitochondrial probe | Identifier | Source |
| --- | --- | --- |
| MitoSOX Mitochondrial Superoxide Indicators, for live-cell imaging | #M36008 | Invitrogen by Thermo Fisher Scientific |
| CM-H2DCFDA (General Oxidative Stress Indicator) | #C6827 |  |
| JC-1 Dye (Mitochondrial Membrane Potential Probe) | #T3168 |  |
| MitoTracker Deep Red FM | #M22426 |  |
| MitoTEMPO | #SML0737 | Sigma-Aldrich |
| Methyl-triphenylphosphoniumchlorid | #468002 |  |

### Antibodies and fluorescent dyes

| Target | Fluorochrome | Clone | Isotype | Host species | Identifier | Source |
| --- | --- | --- | --- | --- | --- | --- |
| CD45 | APC-Cy7 | 30-F11 | IgG<sub>2b</sub>,κ | Rat | #103116 | BioLegend |
| Ly6G | APC | 1A8 | IgG<sub>2a</sub>,κ | Rat | #127614 |  |
| CD11b | FITC | M1/70 | IgG<sub>2b</sub>,κ | Rat | #101206 |  |
| CD11c | PE-Cy7 | N418 | Armenian Hamster IgG | Armenian Hamster | #117318 |  |
| 7-AAD | 7-AAD | n.a. | n.a. | n.a. | #420404 |  |
| Annexin V | Pacific-Blue | n.a. | n.a. | n.a. | #640918 |  |
| CD170/<br>Siglec-F | BV421 | E50-2440 | IgG<sub>2a</sub>,κ | Rat | #565934 | BD Biosciences |
| n.a. | CellTrace™<br>Calcein Red-Orange, AM | n.a. | n.a. | n.a. | #C34851 | Invitrogen |
| Tomm20 | Recombinant<br>Alexa Fluor®<br>555 Anti-TOMM20<br>antibody | EPR15581-54 | Alexa Fluor®<br>555 Rabbit<br>IgG, mono-<br>clonal<br>(EPR25A) | Rabbit | #ab22129<br>2 | Abcam |
| UCP2 | UCP2<br>(D1O5V)<br>Rabbit mAb | D1O5V | Rabbit IgG | Rabbit | #89326 | Cell<br>Signaling |
| β-Actin | β-Actin<br>Antibody | n.a. | n.a. | Rabbit | #4967 |  |

### Critical commercial assays and kits

| Assay | Identifier | Source |
| --- | --- | --- |
| WST-1 Assay Kit (Cell Proliferation) | #ab65473 | Abcam |
| Seahorse XFp Cell Mito Stress Test Kit | #103010-100 | Agilent |
| Seahorse XF Cell Mito Stress Test Kit | #103015-100 |  |
| Seahorse XFp Mito Fuel Flex Test Kit | #103270-100 |  |
| Seahorse XF Mito Fuel Flex Test Kit | #103260-100 |  |
| Bio-Plex Pro Human Cytokine 27-plex Assay | #M500KCAF0Y | BIO-RAD |
| Bio-Plex Pro Mouse Cytokine 23-plex Assay | #M60009RDPD |  |
| RNeasy Micro Kit | #74004 | Qiagen |
| DNeasy Blood & Tissue Kit | #69506 |  |
| Macrophage Isolation Kit (Peritoneum), mouse | #130-110-434 | Miltenyi Biotec |
| MACSxpress Whole Blood Neutrophil Isolation Kit, human | #130-104-434 |  |
| Neutrophil Isolation Kit, mouse | #130-097-658 |  |
| Glucose-Glo | #J6021 | Promega |
| Lactate-Glo | #J5021 |  |
| Glutamine/Glutamate-Glo | #J8021 |  |

### Chemicals, peptides, and recombinant proteins

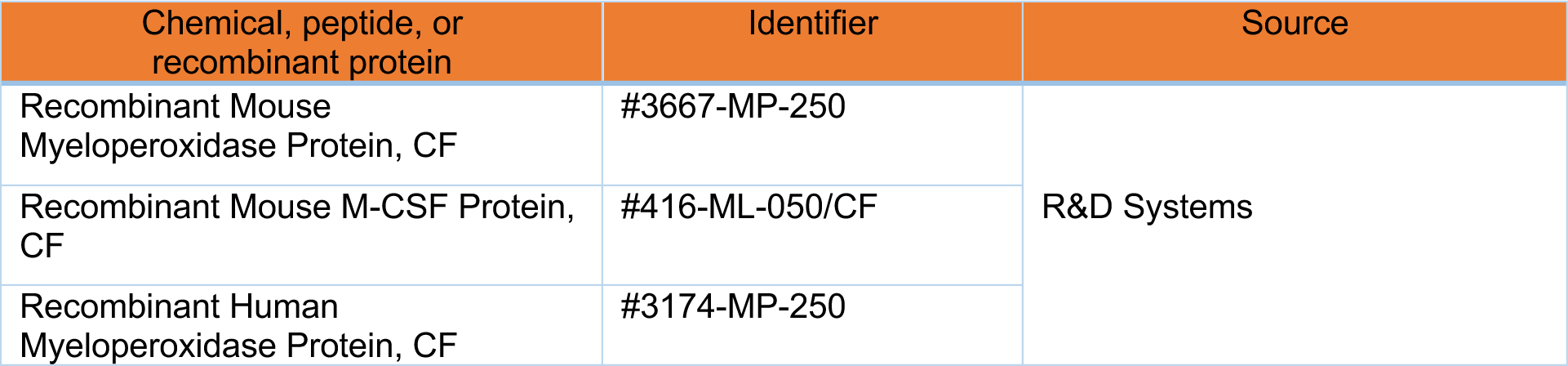
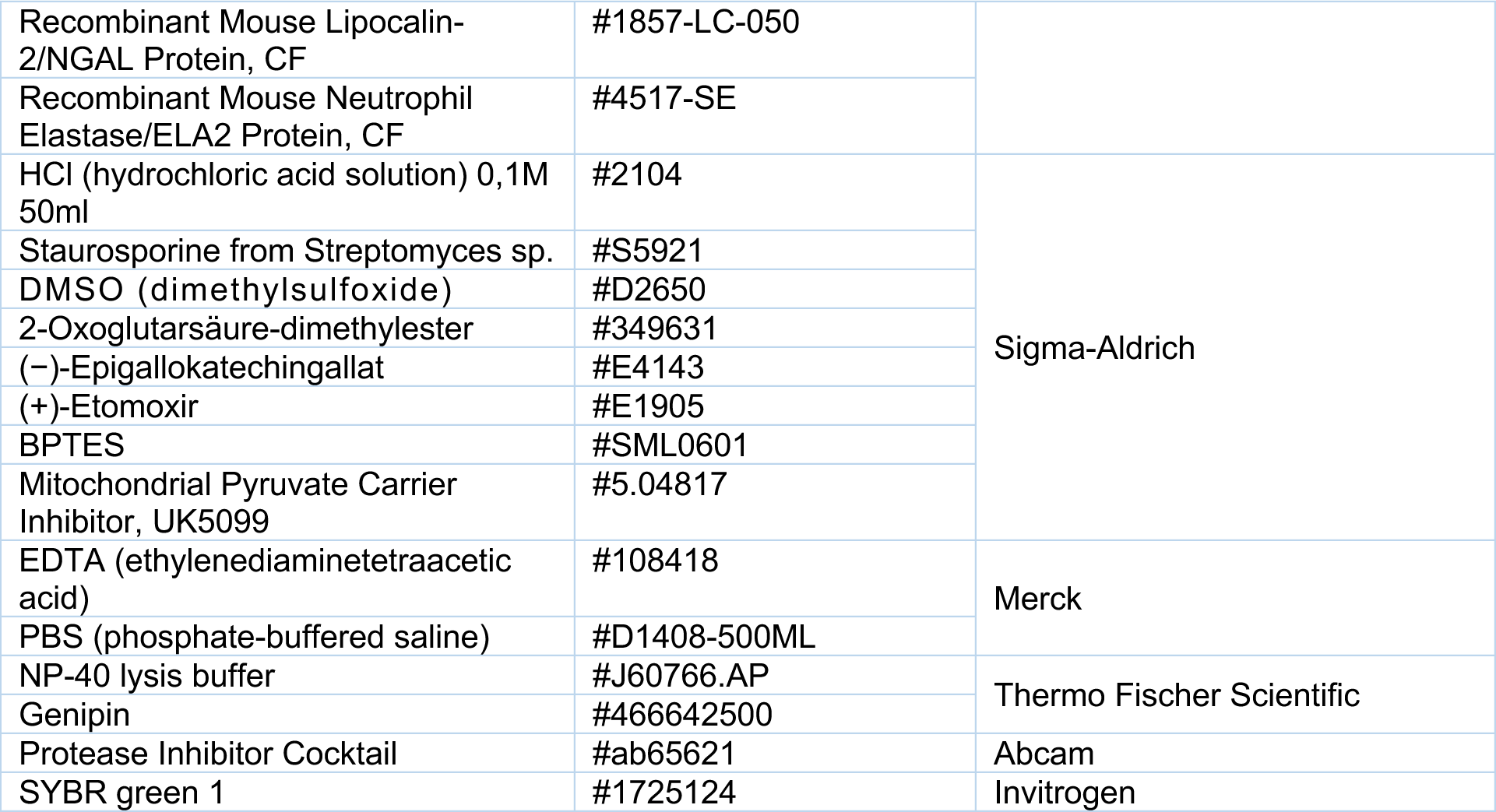

### Cell culture/ *in-vitro* assay media and supply

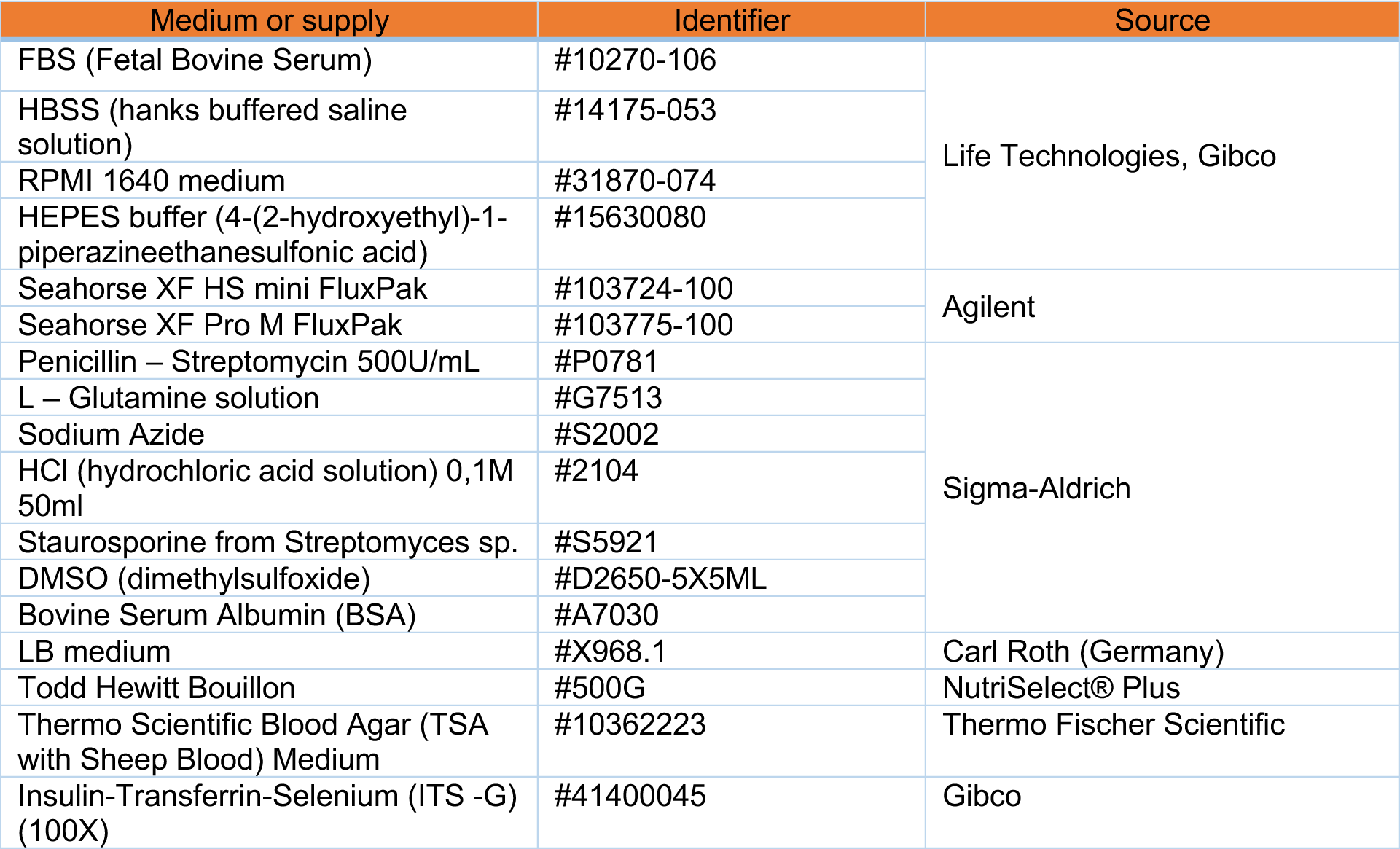

### Anesthesia and in vivo experiments

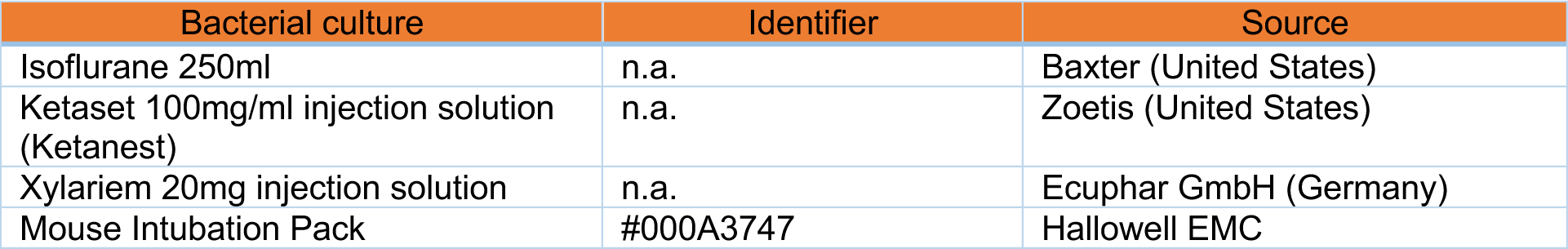

### Bacteria and virus

| Bacterial culture | Identifier | Source |
| --- | --- | --- |
| *K. pneumoniae* | 700721 | American Type Culture Collection |
| *S. pneumoniae* (PN23) | PN23 | Cryo stock |
| *P. aeruginosa* (PA103) | PA103 |  |
| Influenza A Virus | Mouse-adapted PR8 |  |

### Experimental models: Organisms/strains

| Organisms/strains | Genetic Background | Source |
| --- | --- | --- |
| WT | C57BL/6N | Charles River |
| *Ucp2*<sup>-/-</sup> | B6.129S4-Ucp2tm1Lowl | Laboratory N. Sommer, Justus-Liebig-University Gießen, Germany |
| MPO<sup>-/-</sup> | B6.129X1-Mpotm1Lus/J | Charles River |

### Software

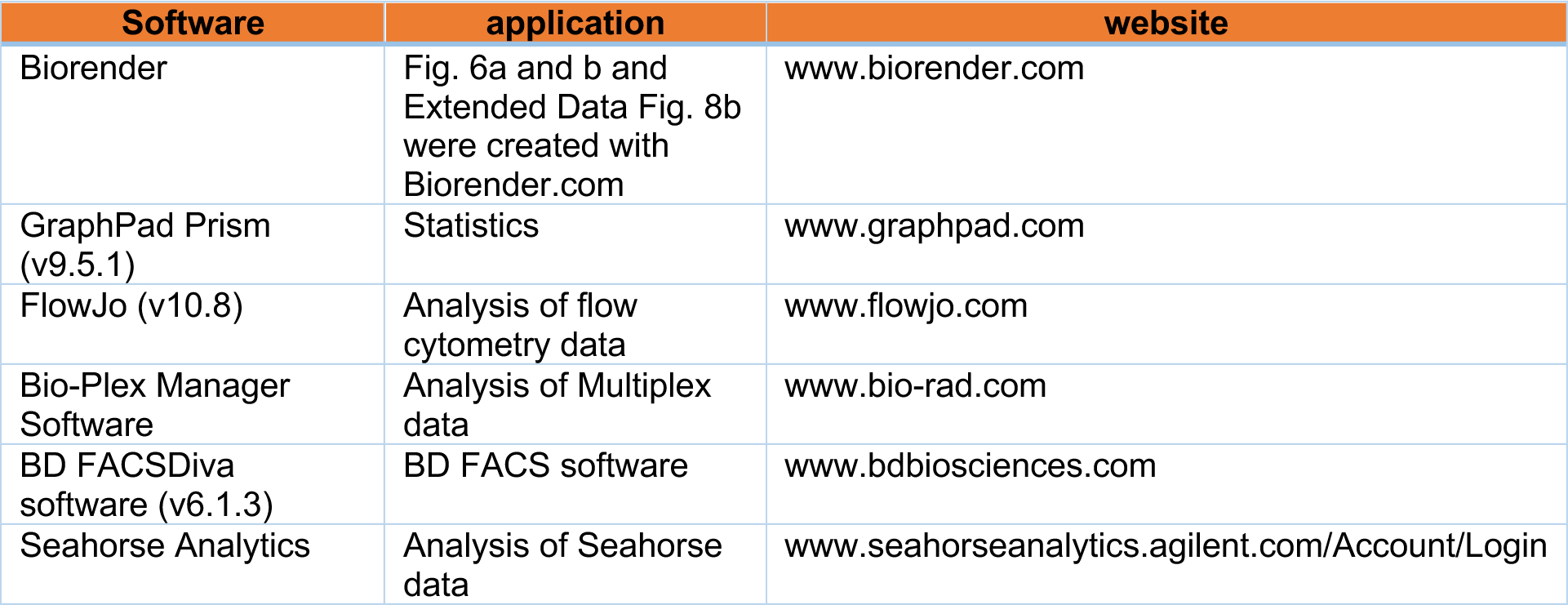

| Software | application | website |
| --- | --- | --- |
| Biorender | Fig. 6a and b and Extended Data Fig. 8b were created with Biorender.com | www.biorender.com |
| GraphPad Prism (v9.5.1) | Statistics | www.graphpad.com |
| FlowJo (v10.8) | Analysis of flow cytometry data | www.flowjo.com |
| Bio-Plex Manager Software | Analysis of Multiplex data | www.bio-rad.com |
| BD FACSDiva software (v6.1.3) | BD FACS software | www.bdbiosciences.com |
| Seahorse Analytics | Analysis of Seahorse data | www.seahorseanalytics.agilent.com/Account/Login |

**Figure S1.**
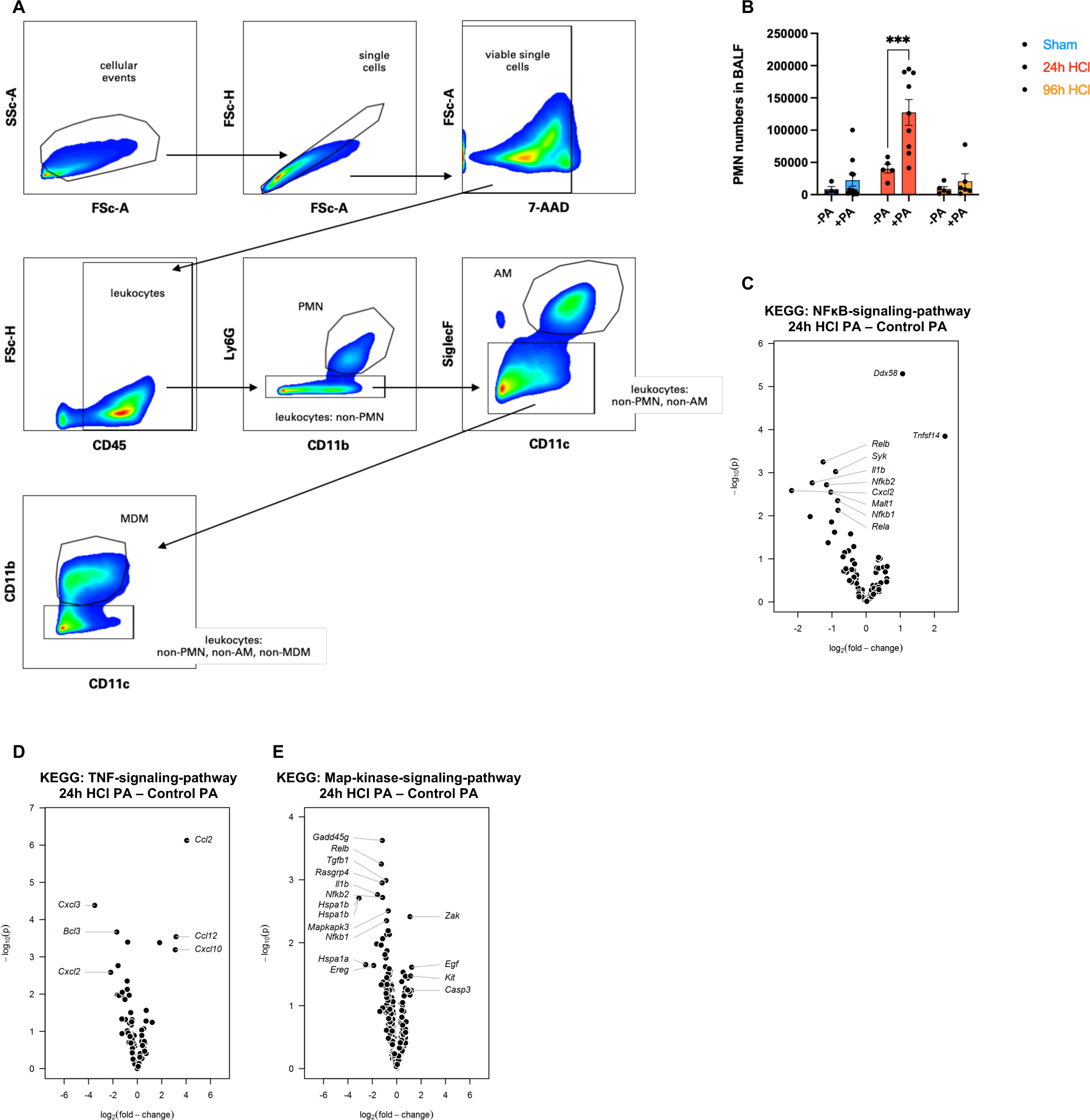
Functional and transcriptional properties of AMs during acid aspiration. (**A**) Gating strategy to identify AMs, MDMs, and PMNs. (**B**) PMN numbers in BALF after acid aspiration (24-96 hours HCl) and subsequent infection with PA *in vivo* for 3 hours compared to acid aspiration alone (n=5-9 mice/group, pooled from two independent experiments). **(C** to **E**) KEGG-pathway analysis of bulk mRNA sequencing data of AMs harvested 3 hours after PA infection *in vivo* versus 24 hours after acid aspiration + 3 hours PA infection *in vivo* for NFκB- (**C**), TNF- (**D**), or Map-kinase-signaling pathway (**E**) (Bulk mRNA sequencing: n=4 mice/group). *p < 0.05, **p < 0.01, ***p < 0.001, ****p < 0.0001 by two-way ANOVA with Sidak’s multiple comparisons test (**B**). Statistical significance in (**C** to **E)** is presented as negative log p-values. Data are shown as mean ± s.e.m.

**Figure S2.**
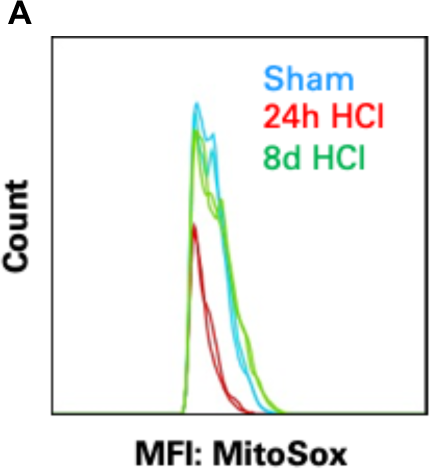
Determining mitochondrial reactive oxygen species production by AMs after acid aspiration. **(A)** Exemplary flow cytometric analysis (histogram) of Fig. 2A depicting the mean fluorescence signal of AMs stained with MitoSox after acid aspiration at indicated time-points in response to *P. aeruginosa ex vivo*.

**Figure S3.**
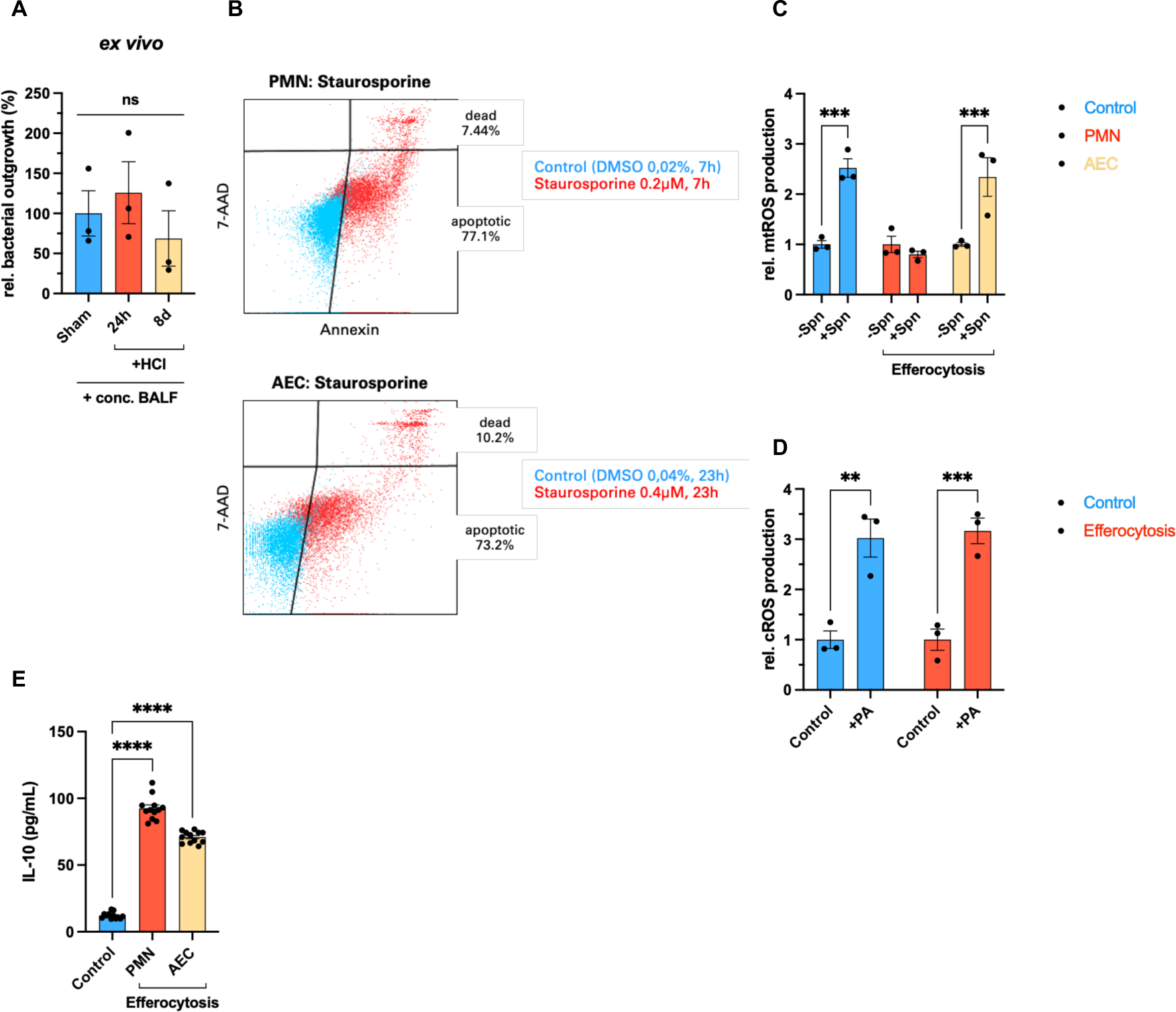
Cell-type specific efferocytosis of PMNs precludes mtROS generation in response to bacteria in AMs. (**A**) Bacterial killing capacity (*P. aeruginosa*) of AMs treated *ex vivo* for 16 hours with concentrated BALF of mice that received HCl intratracheally for 24 hours compared to 8 days or sham treatment (n=3 replicates/group, pooled from four mice/group and two independent experiments). (**B**) Quadrant analysis of annexin V^+^ and propidium iodide^-^ from apoptotic PMNs (top) and AECs (bottom) after incubation with staurosporine 0.2µM for 7 hours or 0.4µM for 23 hours, respectively. (**C**) mtROS production by AMs after clearance of apoptotic PMNs or AECs for 4 hours in response to *S. pneumoniae* for 6-8 hours (n=3 replicates/group, pooled from four mice/group and two independent experiments). (**D**) Assessment of cROS production after efferocytosis of apoptotic PMNs for 4 hours in response to PA for 6-8 hours (n=3 replicates/group, pooled from four mice/group and two independent experiments). (**E**) IL-10 secretion of AMs after efferocytosis of apoptotic PMNs or AECs for 4 hours (n=10-12 replicates/group, pooled from six mice/group and two independent experiments). *p < 0.05, **p < 0.01, ***p < 0.001, ****p < 0.0001 by one-way ANOVA with Tukey’s multiple comparisons test (**A**, **E**) or two-way ANOVA with Sidak’s multiple comparisons test (**C** and **D**). Data are shown as mean ± s.e.m.

**Figure S4.**
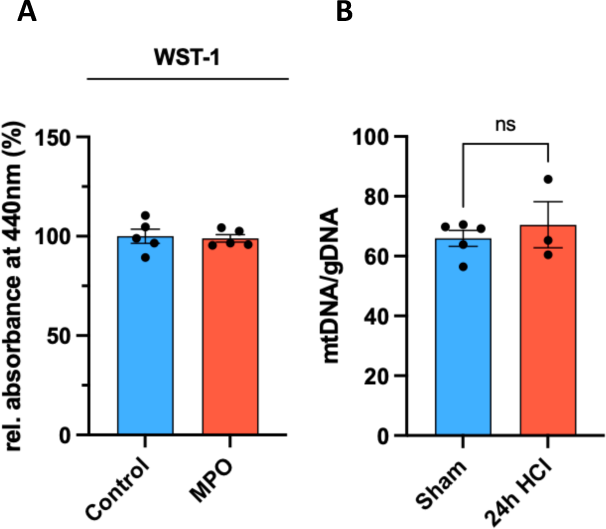
PMN-derived MPO mediates alterations of mitochondrial signaling in AMs. (**A**) Assessment of cell viability of AMs +/− MPO 0,5μM for 3 hours using the WST-1 assay (n=5 replicates/group, pooled from three mice/group and two independent experiments). (**B**) Mitochondrial content 24 hours after acid aspiration compared to sham treatment determined by the ratio of mitochondrial to genomic DNA (mtDNA/gDNA) as measured by quantitative PCR (n=3-5 replicates/group, pooled from 3-4 mice/group and two independent experiments). *p < 0.05, **p < 0.01, ***p < 0.001, ****p < 0.0001 by Student’s t-test (**A** and **B**). Data are shown as mean ± s.e.m.

**Figure S5.**
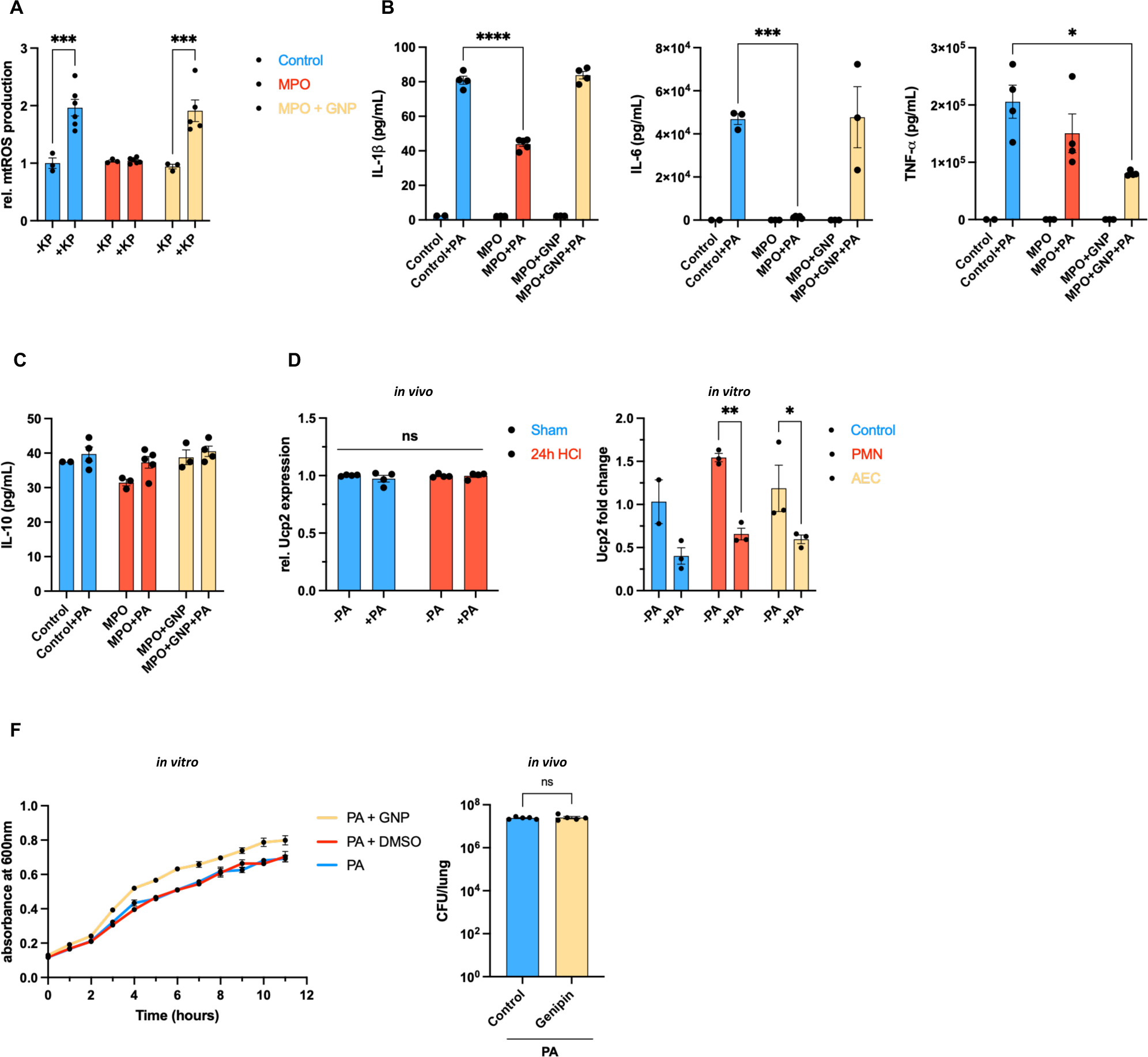
MPO-mediated increase of UCP2 precludes mtROS release in response to bacteria. (**A**) mtROS production in AMs after pre-treatment with MPO 0.5μM for 3 hours +/− GNP 100μM for 1 hour in response to *K. pneumoniae* (KP) for 6-8 hours (n=3-6 replicates/group, pooled from 3-5 mice/group and two independent experiments). (**B**) Release of IL-1β (left), IL-6 (middle), and TNF-α (right) in response to PA after pre-treatment with MPO 0.5μM for 3 hours +/− GNP 100μM for 1 hour (n=3-5 replicates/group, pooled from 3-4 mice/group and two independent experiments). (**C**) Release of IL-10 in response to PA after pre-treatment with MPO 0.5μM for 3 hours +/− GNP 100μM for 1 hour (n=3-5 replicates/group, pooled from 3-4 mice/group and two independent experiments). (**D**) Transcriptional regulation of UCP2 at different time points after acid aspiration + PA for 3 hours *in vivo* (Bulk mRNA sequencing data) (left) and after efferocytosis of apoptotic PMNs or AECs for 4 hours + PA for 6-8 hours *in vitro* (qPCR) (right) (right: n=3 replicates/group, pooled from 5 mice/group and two independent experiments). (**E**) Bacterial growth curves of PA in presence of GNP (100μm), control (DMSO), or untreated (measurement of absorbance at 600nm in technical triplicates every hour for 11 hours) (left); Bacterial burden in lung after intratracheal instillation of Genipin 300μM for 2 hours compared to control (DMSO) and subsequent infection with PA *in vivo* (right) (left: n=3 replicates/group; right: n=5 mice/group). *p < 0.05, **p < 0.01, ***p < 0.001, ****p < 0.0001 by Student’s t-test (**E**) or two-way ANOVA with Sidak’s multiple comparisons test (**A** to **D**). Data are shown as mean ± s.e.m.

**Figure S6.**
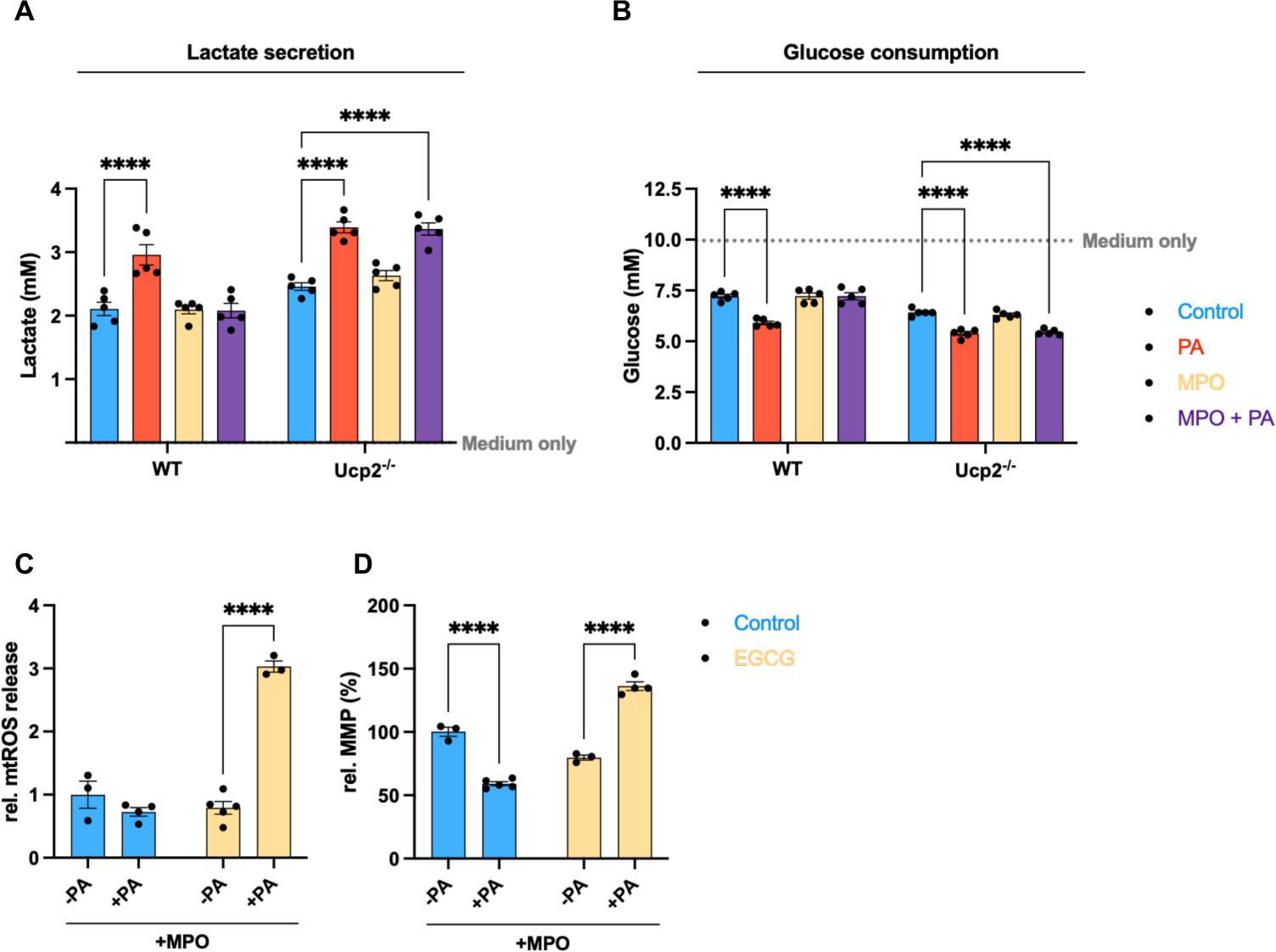
MPO drives canonical glutaminolysis through UCP2. (**A** and **B**) Lactate secretion (A) and glucose consumption (B) by AMs from WT or *Ucp2^−/−^* mice after pre-treatment with MPO 0.5µm for 3 hours in response to PA for 6-8 hours determined by Luminometry (n=5 replicates/group, pooled from 4 mice/group and two independent experiments). (**C**) mtROS production by AMs after pre-treatment with MPO 0.5µm for 3 hours +/− EGCG 100µm for 1 hour in response to *P. aeruginosa* (n=3-5 replicates/group, pooled from 3-4 mice/group and two independent experiments). (**D**) MMP of AMs after pre-treatment with MPO 0.5µm for 3 hours +/− EGCG 100µm for 1 hour in response to PA (n=3-5 replicates/group, pooled from 3-4 mice/group and two independent experiments). *p < 0.05, **p < 0.01, ***p < 0.001, ****p < 0.0001 by two-way (**C** and **D**) or three-way ANOVA (**A** and **B**) with Sidak’s multiple comparisons test. Data are shown as mean ± s.e.m.

**Figure S7.**
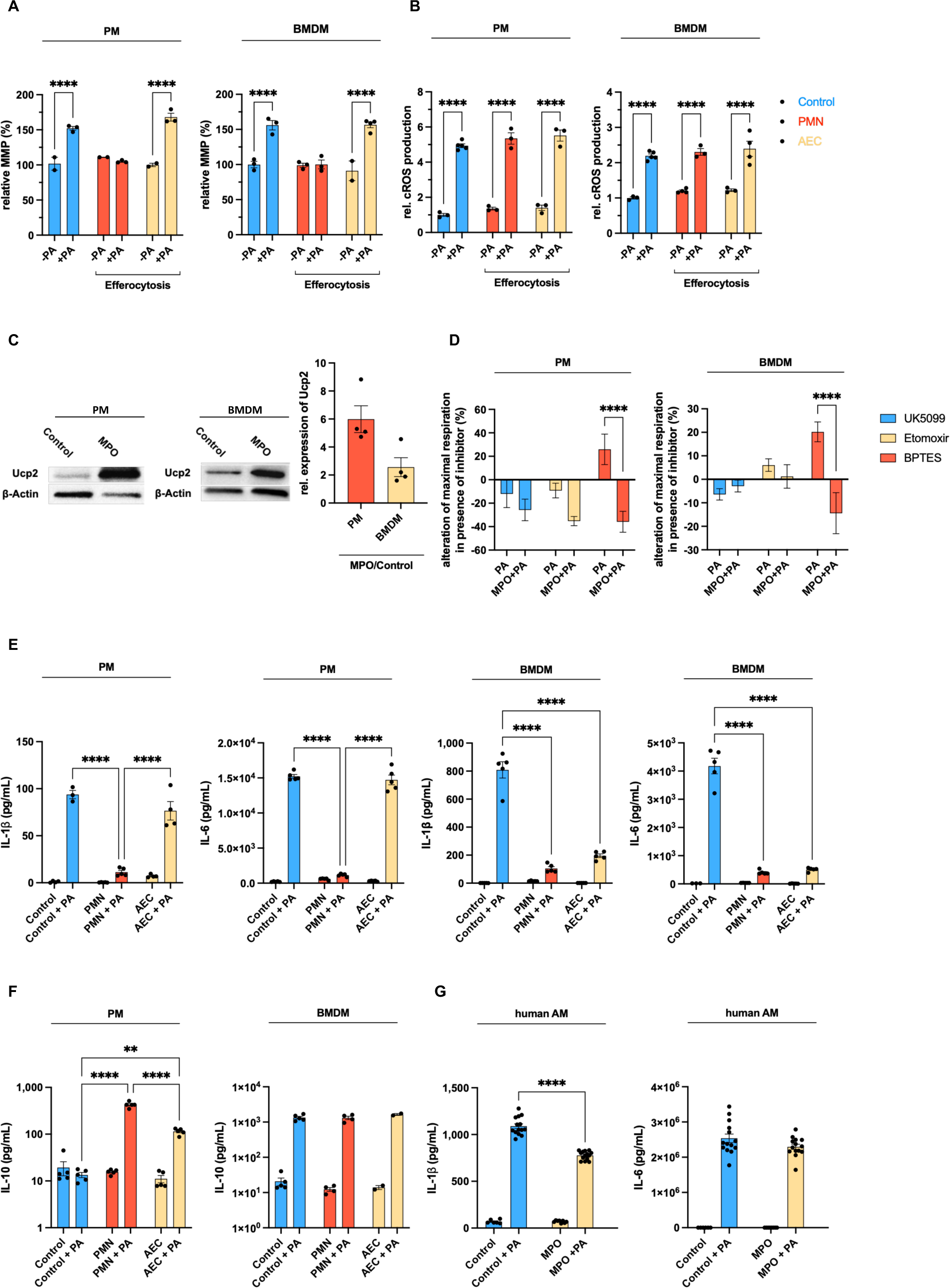
MPO acts via UCP2 to impair bacterial control in different macrophages across species. (**A**) Assessment of MMP in PMs (left) and BMDMs (right) after pre-treatment with MPO 0.5µm for 3 hours in response to PA for 6-8 hours (n=3-4 replicates/group, pooled from 3-4 mice/group and two independent experiments). (**B**) Assessment of cROS in PMs (left) and BMDMs (right) after pre-treatment with MPO 0.5µm for 3 hours in response to PA for 6-8 hours (n=3-5 replicates/group, pooled from 3-4 mice/group and two independent experiments). (**C**) Western Blot of UCP2 and β-Actin (loading control) in PMs (left) and BMDMs (right) +/− MPO 0.5µm for 3 hours (n=4 replicates/group, pooled analysis from 8-11 mice/group and two independent experiments). (**D**) Substrate Oxidation Stress test: Alteration of maximal respiration in presence of pathway-specific inhibitors (UK5099 for glycolysis, etomoxir for fatty acid oxidation, BPTES for glutaminolysis) compared to respective uninhibited control in PMs (left) and BMDMs (right) treated with MPO 0.5µM for 3 hours in response to PA for 6-8 hours (n=9-10 replicates/group, pooled from 8-10 mice/group and two independent experiments). (**E**) IL-1β and IL-6 secretion of PMs (left) and BMDMs (right) after efferocytosis of apoptotic PMNs for 4 hours in response to PA (n=3-5 replicates/group, pooled from 3-4 mice/group and two independent experiments). (**F**) IL-10 secretion of PMs (left) and BMDMs (right) after efferocytosis of apoptotic PMNs for 4 hours in response to PA (n=3-5 replicates/group, pooled from 3-4 mice/group and two independent experiments). (**G**) Secretion of IL-1β and IL-6 of human AMs after pre-treatment with MPO 0.5µM for 3 hours in response to PA (n=6-14 replicates/group, pooled from three patients and three independent experiments). *p < 0.05, **p < 0.01, ***p < 0.001, ****p < 0.0001 by two-way ANOVA with Sidak’s multiple comparisons test (**A** and **B, D** to **G**). Data are shown as mean ± s.e.m.

**Figure S8.**
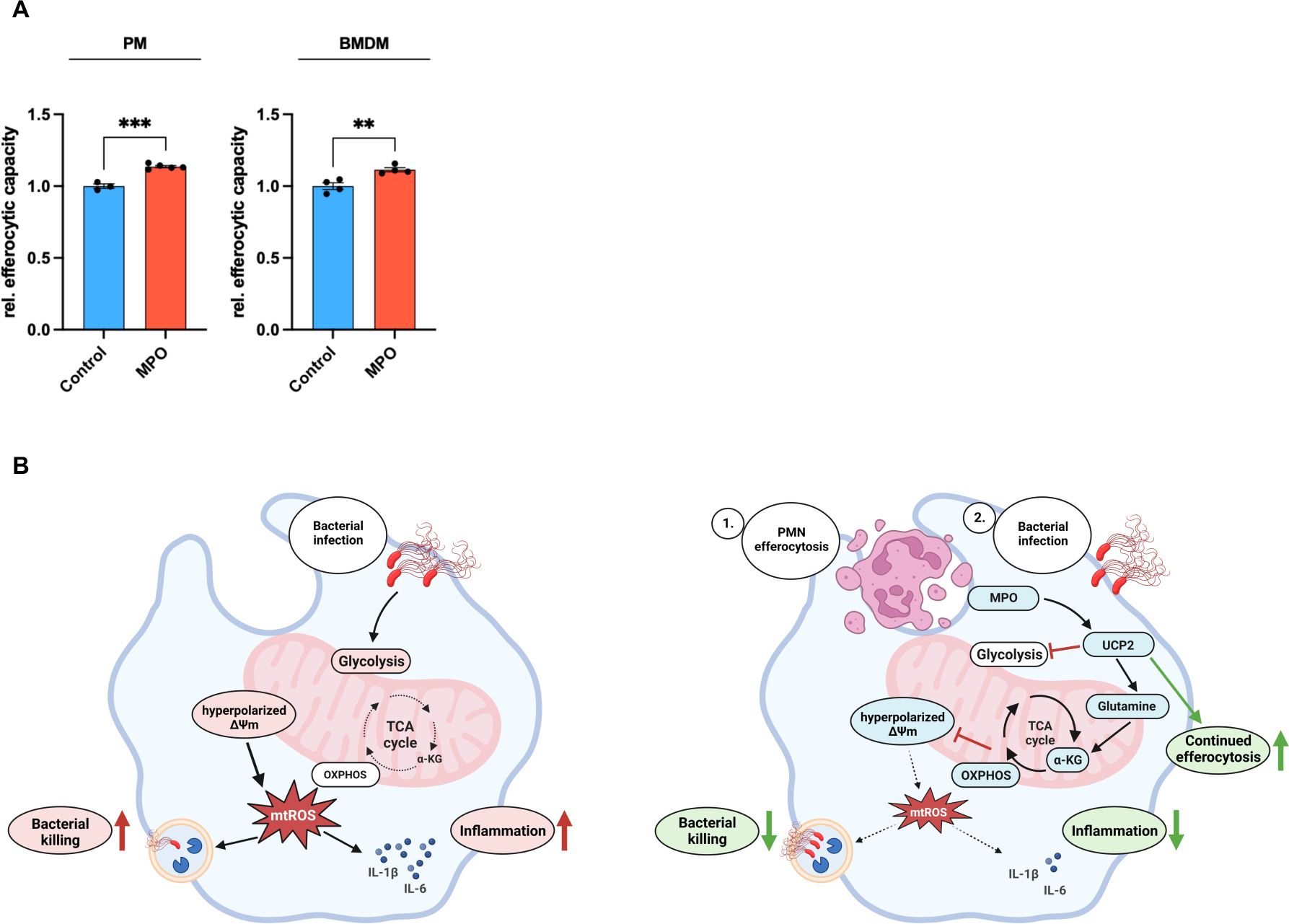
MPO acts via UCP2 to improve efferocytic capacity in different macrophages. (**A**) Efferocytic capacity of PMs (left) and BMDMs (right). PMs and BMDMs were treated with MPO 0.5µM for 3 hours compared to control before incubating all samples with pre-stained apoptotic PMNs for 1 hour. Efferocytic capacity was then quantified by flow cytometry (n=3-5 replicates/group, pooled from 3-4 mice/group and two independent experiments). *p < 0.05, **p < 0.01, ***p < 0.001, ****p < 0.0001 by Student’s t-test. Data are shown as mean ± s.e.m. (**B**) Graphical representation of proposed mechanism.

